# Single-cell analysis of the muscle stem cell hierarchy identifies heterotypic communication signals involved in skeletal muscle regeneration

**DOI:** 10.1101/671032

**Authors:** Andrea J. De Micheli, Paula Fraczek, Sharon Soueid-Baumgarten, Hiranmayi Ravichandran, Iwijn De Vlaminck, Olivier Elemento, Benjamin D. Cosgrove

## Abstract

Muscle stem cells (MuSCs) are an essential adult stem cell population with the capacity to self-renew and regenerate muscle tissue. Functionally heterogeneous subpopulations of MuSCs have been identified based on their expression of myogenic regulatory factors and surface markers. However, a unified organization of muscle stem and progenitor cells and their subpopulations remains unresolved. Here, we performed temporal analysis of skeletal muscle regeneration using single-cell RNA-sequencing (scRNA-seq) of myotoxin-injured adult mouse hindlimb muscles. We generated over 34,000 single-cell transcriptomes spanning four muscle regeneration time-points and identified 15 distinct cell types, including a heterogeneous population of MuSCs and progenitor cells. Our analysis provides a hierarchical map of myogenic cell populations and identifies stage-specific regulatory programs that govern their contributions to muscle regeneration. In this transcriptomic atlas, we observed cell type-specific regenerative dynamics, exemplified by waves of transient amplification and diversification of multiple immune cell types and, subsequently, myogenic cells. Unbiased trajectory inference organized the myogenic cell populations within the atlas into a continuum, consisting of a hierarchy of quiescent MuSCs, cycling progenitors, committed myoblasts, and terminally differentiated myocytes. This myogenic trajectory matched prior understanding and also revealed that MuSC stages are defined by synchronous changes in regulatory factors, cell cycle-associated, and surface receptor gene expression. Lastly, we analyzed the transcriptomic atlas to identify over 100 candidate heterotypic communication signals between myogenic and non-myogenic cell populations, including many involving the fibroblast growth factor (FGF), Notch, and Syndecan receptor families and their associated ligands. Syndecan receptors were implicated in a large fraction of these cell communication interactions and were observed to exhibit transcriptional heterogeneity within the myogenic continuum. Using multiparameter mass cytometry (CyTOF), we confirmed that cycling MuSCs exhibit diversified Syndecan-1/2 expression, which suggests that dynamic alterations in Syndecan signaling interactions may coordinate stage-specific myogenic cell fate regulation. This scRNA-seq reference atlas provides a resolved hierarchical organization of myogenic subpopulations as a resource to investigate cell-cell interactions that regulate myogenic stem and progenitor cell fates in muscle regeneration.

## Introduction

Muscle stem cells (MuSCs), also known as satellite cells, are essential for skeletal muscle homeostasis and regeneration throughout lifespan (Blau et al., 2015; Wang and Rudnicki, 2011). MuSCs are found at the periphery of the muscle myofibers and are sheltered inside a niche microenvironment where they are maintained in a quiescent state. In response to injury, MuSCs activate, self-renew, and differentiate into progenitors capable of myofiber repair. This regenerative process is orchestrated by a network of interactions with a variety of cell types including immune cells, endothelial cells, and fibro/adipogenic progenitors (FAPs) (Wosczyna and Rando, 2018). For example, FAPs secrete fibronectin, insulin-like growth factor-1, and other matrix proteins and growth factors to coordinate muscle tissue repair through the regulation of myogenic cell fates and the clearance of cellular debris (Heredia et al., 2013; Joe et al., 2010; Lukjanenko et al., 2016).

A continuum of myogenic stem and progenitor cell populations are present in regenerating muscle (Motohashi and Asakura, 2014; Tierney and Sacco, 2016). MuSCs are quiescent Pax7-expressing cells in homeostasis and, following injury, progress in the cell cycle and exhibit an activated myogenic expression program marked by expression of Myf5 (Wang and Rudnicki, 2011). Following cell division, their progeny either self-renew to replenish the Pax7^+^ MuSC pool or differentiate into MyoD^+^ myogenic progenitors (myoblasts), which later commit to fusion-competent Myogenin^+^ myocytes. This concept of the myogenic cell lineage was largely derived from lineage tracing and prospective isolation studies using myogenic regulatory factors and cell cycle to define cell states (Biressi and Rando, 2010). Myogenic stem/progenitor cells populations, purified through surface antigen profiles and/or transgenic reporters, exhibit substantial molecular and functional heterogeneity throughout adulthood (Chakkalakal et al., 2014; Cornelison and Wold, 1997; Cosgrove et al., 2014; Kuang et al., 2007; Porpiglia et al., 2017; Rocheteau et al., 2012; Sacco et al., 2008; Sousa-Victor et al., 2014; Tierney et al., 2018). These findings suggest that myogenic stem/progenitor cell lineage can be interpreted as a continuum of hierarchical cell states. However, it remains an unresolved challenge how global profiles in cell cycle mediators, regulatory factors and surface markers define this myogenic continuum.

Recent advances in single-cell analyses and algorithms provide potent new strategies to infer cell differentiation trajectories (Hwang et al., 2018; Wagner et al., 2016). Here, we generated a single-cell transcriptomic atlas of mouse muscle regeneration to describe the myogenic continuum and multicellular communication networks involved in muscle repair in mice. We used droplet-based single-cell RNA-sequencing (scRNA-seq) to collect a multi-cell-type transcriptomic reference time-course, spanning four time-points and over single-cell 34,000 transcriptomes, of the regenerating muscle tissue in mice. We analyze this atlas to identify the compositional and gene expression dynamics of the cellular constituents of muscle repair in a manner not biased by prior knowledge or lineage tracing. Using trajectory inference, we organized more than 3,200 individual myogenic cell transcriptomes in a pseudo-time continuum to reveal their hierarchical organization and identify regulatory factor and surface marker expression profiles unique to distinct myogenic subpopulations. Finally, we used a ligand-receptor synchronous coexpression model to identify cell communication interactions between non-myogenic cells and distinct myogenic subpopulations involved the muscle repair process.

## Results

### A single-cell RNA-sequencing atlas of muscle regeneration

Skeletal muscle regeneration in response to local tissue damage depends on the coordinated interactions of multiple myogenic and non-myogenic cell types over a time-course of weeks (Wosczyna and Rando, 2018). To gather a comprehensive view of this process, we generated transcriptomic atlas of adult mouse hindlimb muscle regeneration using droplet-based single-cell 3’ RNA-sequencing (scRNA-seq) on the 10X Chromium platform. We collected tibialis anterior (TA) muscles of healthy adult (4-6 month) C57BL6 mice at 0, 2, 5, and 7 days following injection of the myotoxin notexin to induce myofiber damage (n=2-3 mice per time-point). We dissected and then enzymatically digested the TA muscles into single-cell suspensions and then filtered to remove cellular debris and, in some samples, applied red blood cells (RBC) lysis before performing scRNA-seq (**Fig. 1A**).

**Figure 1.**
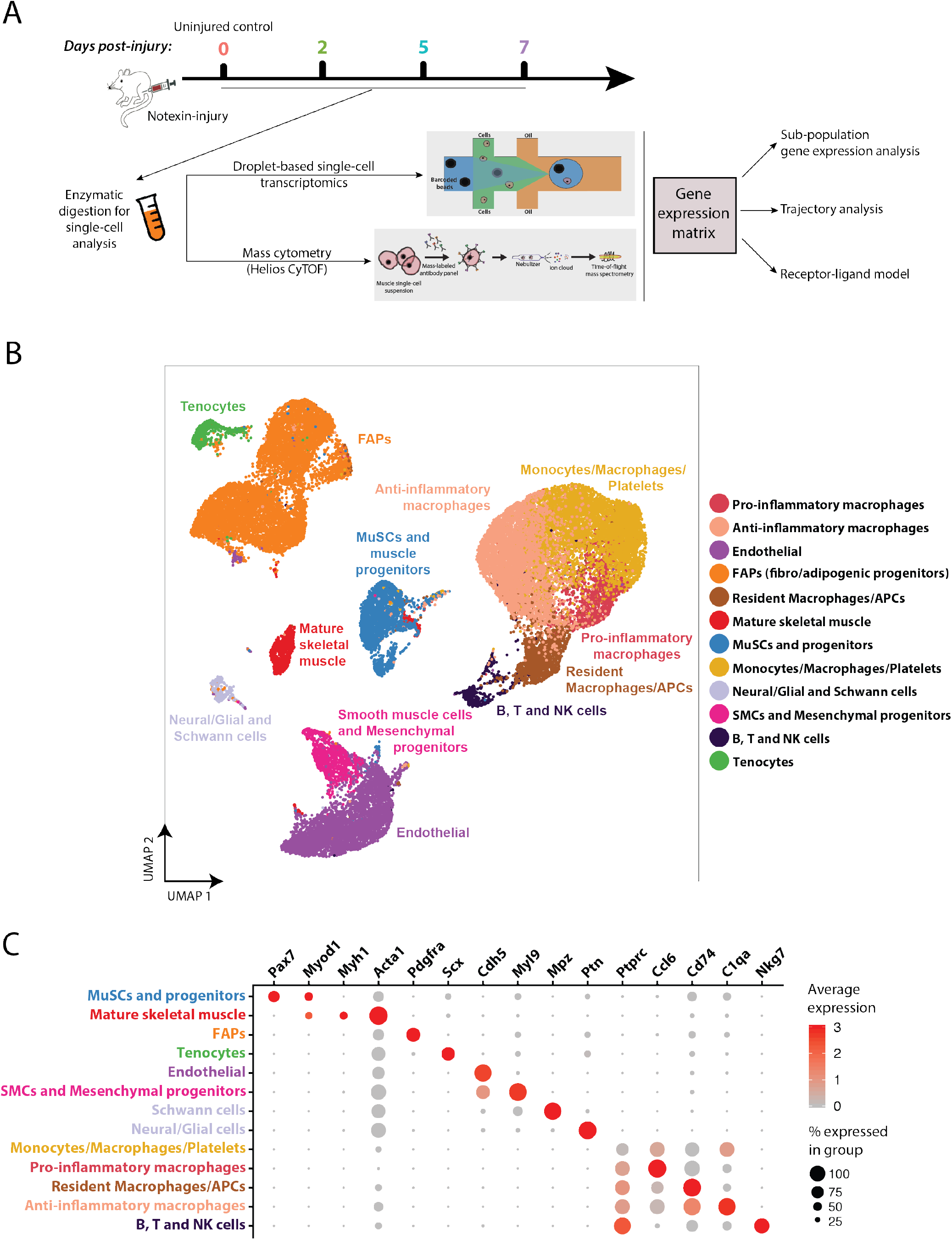
Assembly and curation of a scRNA-seq atlas of muscle regeneration. (**A**) Experimental design overview. Cell suspensions were collected from digested tibialis anterior (TA) muscles of adult mice at various time points (0, 2, 5, and 7 days) following notexin injury (n=2-3) and subjected to scRNA-seq and mass cytometry (CyTOF), followed by downstream analyses. (**B**) Complete 34,438 cell transcriptomic atlas assembled from all sample time-points. Data is presented as a UMAP projection used to visualize variation in singlecell transcriptomes. Unsupervised SNN clustering resolved at least 12 distinct types of cells (color-coded in legend). More resolved cell type clusters, distinguishing Neural/Glial from Schwann cells, immature B from cytotoxic T cells, were evident when analyzing time-points individually (see **Fig. 2A**). (**C**) Identification of cell types from SNN clusters based on cluster-average expression of canonical genes. Dot size represents the percentage of cells with a nonzero expression level and color-scale represents the average expression level across all cells within cluster.

We used the Seurat package for scRNA-seq data filtering and processing (see **Methods**). We removed cells with fewer than 200 genes detected, fewer than 1000 UMIs, or with more than 20% of UMIs mapped to mitochondrial genes (**Fig. S1-1A-B**). Applying these filters eliminated erythroblasts to levels similar to those observed in samples where RBC lysis was performed, suggesting that the RBC step could be omitted (**Fig. S1-1D**). We found that the use of Calcein-AM based FACS sorting to enrich for viable cells altered the cellular composition. Specifically, cell sorting based on Calcein-AM positivity decreased the incidence of myogenic (MuSCs and progenitors) and anti-inflammatory macrophage populations (**Fig. S1-2**). This is could be due to biases introduced by viability selection, perhaps owing to reduced metabolism of Calcein-AM by quiescent cells. Based on this observation, we omitted FACS viability sorting in the collection of these scRNA-seq data.

After filtering, the scRNA-seq datasets each contained on average 3,444 ± 1,286 cells. We compared time-point replicate datasets generated from different mice (n=2-3) to evaluate batch effects and mouse-to-mouse variability. We observed only minor differences, most notably a small variation in the incidence of macrophage population in day 7 post-injury samples (**Fig. S1-1C**), suggesting that these datasets contained minimal batch effects. Thus, for subsequent analyses, we combined time-point biological replicates, without batch correction, to improve cell sample size and statistical power. Initially, the samples were assembled into a unified transcriptomic atlas containing 34,438 cells, expressing a total of 19,584 detectable genes (**Figs. 1B** and **S1-1A**). We next annotated the cell types involved in this muscle regeneration atlas. We employed uniform manifold approximation and projection (UMAP) to visualize the individual transcriptomes of all cells in the unified dataset (**Fig. 1B**) (Becht et al., 2018). Independently, we performed unsupervised shared nearest neighbor (SNN) clustering, which partitioned cells into 12 groups based on their transcriptomic programs (**Fig. 1B**). We observed more refined groups, potentially revealing additional subtypes and cell states, when SNN clustering was applied to samples collected at individual time-points (see **Fig. 2**). In examining the full atlas, we interpreted the clusters as broadly defining 12 different cell populations. To identify these populations, we examined the normalized expression level and frequency of canonical cell type genes and named them based on their exclusivity in these expression patterns (**Fig. 1C**). To discriminate more ambiguous populations, especially in the immune subpopulation, we performed differential expression analysis using a negative binomial model between cells within the cluster and all other cells in the atlas (**Figs. S1-3** and **S2-1**).

**Figure 2.**
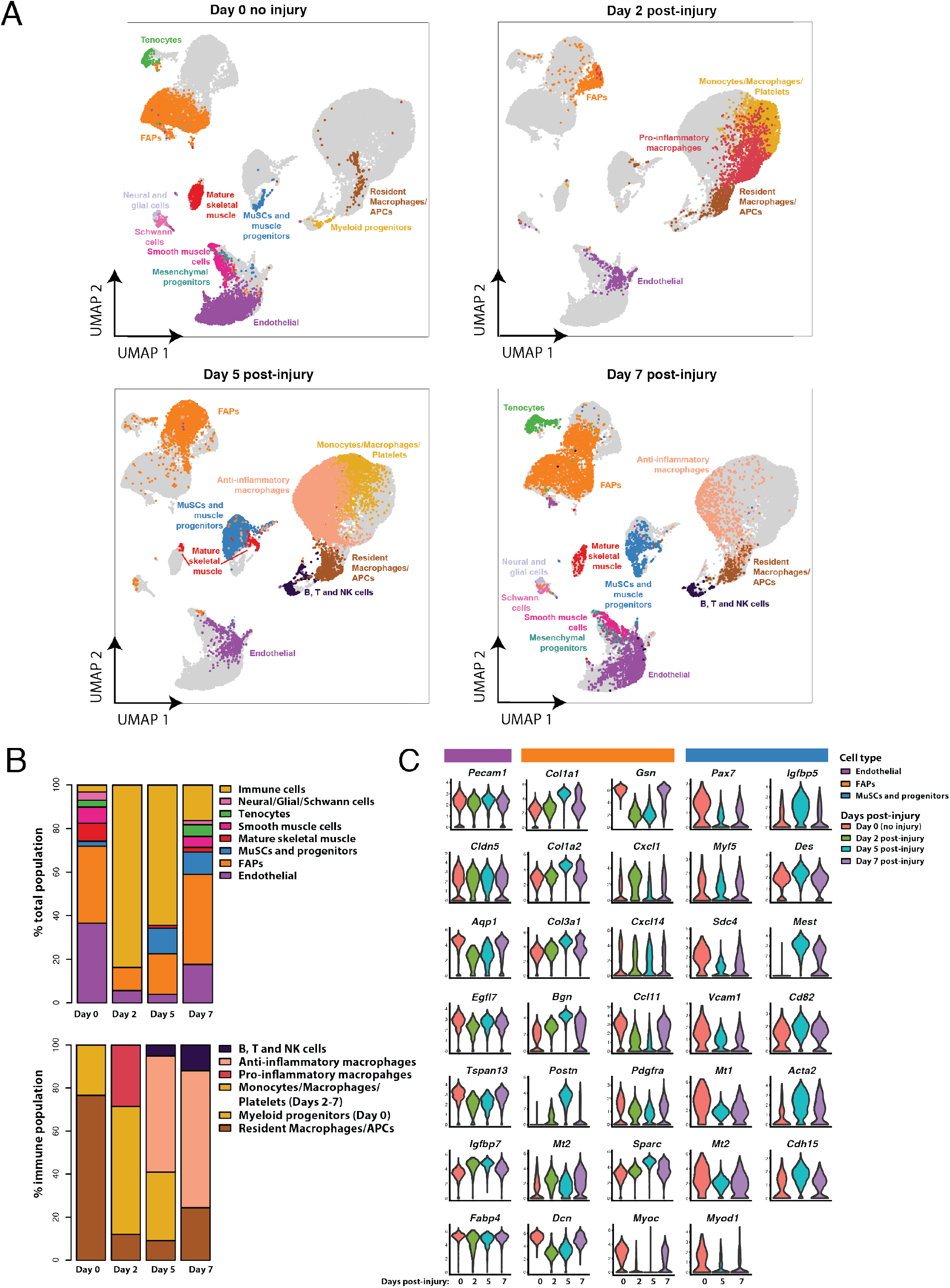
Cell composition and gene expression dynamics of muscle regeneration. (**A**) UMAP projections of muscle single-cell transcriptomes split by time-points post-injury containing, respectively, 7,025, 5,524, 14,240, and 7,646 cells for day 0, 2, 5, and 7 days post-notexin injury. Fourteen total cell types were identified using SNN clustering applied to each time-point. Cells from other time-points are in gray. (**B**) Compositional dynamics of cell types throughout the regeneration time course. Immune cells are grouped together (top) or separated (bottom). (**C**) Violin plots presenting the heterogeneous gene expression changes for a selection of differentially expressed genes within the endothelial, FAP, and MuSC/myogenic progenitors populations at each time-point. MuSCs and myogenic progenitor cells were too rare at day 2 to analyze.

We observed a population of myogenic progenitors, containing MuSCs and myoblasts, which expressed the myogenic transcription factors *Pax7* and *Myod1* (Wang and Rudnicki, 2011). We detected a population of mature myocytes and/or myofibers, which expressed *Myh1* (myosin heavy chain 1) and *Acta1* (skeletal muscle alpha actin 1), both proteins involved in the contractile function of terminally differentiated skeletal muscle cells (Lyons et al., 1990). The limited incidence of this population is likely due to the size-filtering out of multinucleated myofibers during the cell isolation protocol. *Acta1* is also expressed, but at a lower level and frequency, in other cell types, which is not surprising given that it is a common component of the cytoskeleton and plays a role in cell migration. We also identified a population of *Pdgfra*-expressing fibro/adipogenic progenitor cells (FAPs) (Uezumi et al., 2011) and tendon progenitor cells (tenocytes), which express the tenogenic transcription factor *Scx* (scleraxis), and glycoprotein encoding genes *Fmod* (fibromodulin) and *Tnmd* (tenomodulin) (see **Fig. S2**) (Docheva et al., 2005; Schweitzer et al., 2001). Two closely related *Pecam1* (cell adhesion molecule CD31)-expressing populations were identified as, first, smooth muscle cells (SMCs) and mesenchymal progenitors that exclusively express *Myl9* (myosin light chain 9) (Gaylinn et al., 1989) and, second, endothelial cells that express *Cdh5* (cadherin 5) (Christov et al., 2007; Zordan et al., 2014). We also find a mixed group of neuro-muscular cells, which are enriched for expression of the Schwann cell marker *Mpz* (myelin protein zero) and the neuronal gene *Ptn* (pleiotrophin) (Liu et al., 2015).

The cumulatively largest and most ambiguous group are immune cells, which dynamically infiltrate muscle and mediate inflammatory regulation of tissue repair. We observed substantial overlap in immune cell marker gene expression (**Fig. 1C**), which agrees with the immune cell phenotype continuum model (Novak and Koh, 2013). We therefore examined multiple gene signatures to subdivide immune cell clusters (**Figs. S1-4** and **S2-1**) based on prior literature (Chazaud, 2016; Tidball, 2017). One immune cluster contains resident macrophages and antigen presenting cells (APCs), as identified by expression of *Cd74* and other MHCII complex encoding genes. A second immune cluster contains a general group of monocytes, macrophages, and platelets, marked by expression of *Cd68* (common immune glycoprotein CD68) and *Pf4* (platelet factor 4). A third immune cluster contains activated (pro-inflammatory) macrophages as identified by expression of the inflammatory neutrophil and macrophage markers including *Ccl6* (chemokine C-C motif ligand 6). A fourth immune cluster contains antiinflammatory macrophages, as identified by expression of *C1qa*, which encodes for part of the C1 complement complex. A fifth immune cluster contains immature and mature B- and T-lymphocytes and NK cells, as identified by expression of the common lymphocyte gene *Ptprc* (protein tyrosine phosphatase receptor type C, or CD45) and *Nkg7* (natural killer cell granule protein 7). This annotated scRNA-seq dataset provides a reference atlas to examine the cell populations and gene expression dynamics during muscle regeneration.

### Cell type and gene expression dynamics of muscle regeneration at single-cell resolution

Next, we analyzed the transcriptomic atlas to assess how these populations are dynamically altered in incidence and gene expression activity following muscle injury. Overall, we discerned between 5-12 distinct cell populations via SNN clustering at each time-point (**Fig. 2A-B**). Uninjured muscles (day 0) were largely composed of endothelial cells, FAPs, and mature myocytes/myonuclei, with other cell populations detected infrequently. After injury, we observed a transient increase in multiple immune cell types and a concomitant decline in non-immune populations. By day 5, the immune cell population diversified while the myogenic populations recovered. By day 7, most cell populations approached their day 0 frequencies and gene expression states (**Figs. 2C**, **S2-1**, and **S2-2**), revealing a return to homeostasis.

*Pecam1*-expressing endothelial cells were the most abundant cell population at day 0 (38%) and exhibited only minor changes in detection frequency and gene expression heterogeneity throughout the time-course (Christov et al., 2007). In contrast, *Pdgfra*-expressing FAPs exhibited compositional and expression variability. FAPs were a substantial cell fraction of uninjured muscles (31%) and expressed the ECM genes *Col3a1, Dcn*, and *Gsn* in homeostasis (**Fig. 2A-C**, **S2-1**, and **S2-2**). At day 2 post-injury, FAPs also express *Ccl7, Cxcl5*, and *Cxcl1*, chemokines that attract monocytes and neutrophils. FAPs have elevated expression of *Col1a1, Col1a2* and *Mt* and diminished expression of *Gsn* and *Dcn*, suggesting an “activated” or “remodeling” state (**Fig. S2-2**). At day 5, FAPs have further elevated expression of *Col1a1* and *Col1a2* and also express *Postn, Bgn*, and *Sparc* (**Fig. 2C**). At day 7, the FAP expression profile resembles the uninjured samples, albeit with more heterogeneity, indicating a resolution to homeostasis (**Fig. S2-2**).

Likewise, the immune cell populations within this atlas exhibit notable dynamics. At day 0, we detect that immune cell populations comprise 5% of the uninjured muscle, and can be subdivided into a more defined group of *Cd79*^+^ *Ly6d*^+^ immature myeloblasts, *Cd3*^+^ T-cells, and a group of resident macrophages and APCs expressing *Lyz2, Cd74*, and *Ccl6* (**Fig. S1-2, S1-3**). At day 2, the inflammatory response initiates, as reflected by a dramatic increase in the frequency (84%) of immune cells detected (**Fig. 2A**). This early-response immune compartment can be divided into three populations. First, we observed APCs characterized by expression of MHC class II proteins such as *Cd74* and the *H2* family. Within this group we distinguish small populations of *Cd7*^+^ mature T-cells, *Klrd1*^+^ NK cells, and *Cd209a*^+^ dendritic cells. Second, we identified pro-inflammatory macrophages (including activated M1-like macrophages) that express *Ccl9* (a chemokine that attracts *Cd11b*^+^ *Ccr1*^+^ dendritic cells), *Ccr2* (a chemokine involved in monocyte chemotaxis), and *Ly6c2*. Third, we observed a less defined group of cells that express a wide variety of markers including *Cd68*^+^ monocytes and *Pf4*^+^ platelets (**Fig. S1-4, S2-1**). At day 5 post-injury, immune cells still are remain prevalent (64%) (**Fig. 2A**). We observed however a shift in the macrophage population from a pro-inflammatory to an anti-inflammatory phenotype. We found that these macrophages express *C1q* complement genes as well as *Apoe*, characteristic of the M2-like anti-inflammatory phenotype (Baitsch et al., 2011; Ho et al., 2016). We detect some *Aif1*^+^ macrophages and/or dendritic cells that have been reported to modulate muscle repair (Kuschel et al., 2000). We found elevated frequencies of APCs, *Lsp1^+^ Ccr7*^+^ B-cells, *Ccl5*^+^ *Xcl1*^+^ T-cells and *Nkg7*^+^ *Klrd1*^+^ NK cells. (**Fig. S2-1**). At day 7, immune cells were more infrequent (17%) and started to resemble their day 0 cell type composition, with some M2-like macrophages, T-lymphocyte and NK-cells remaining.

The myogenic cell populations exhibited a temporal profile inverted in abundance relative to the immune cells and transcriptionally heterogeneous. At day 0, we detected a small population (2%) of MuSCs and progenitor cells expressing *Pax7, Sdc4, Vcam1*, and *Myod1*, and also detected *Acta1*^+^ *Myl1*^+^ mature myocytes and/or myofiber nuclei (9%). At day 2, we did not detect any MuSCs or progenitors though they are reported to be in an activated state following notexin injury. This surprising finding is likely due the relative rarity of myogenic cells compared immune cells providing a sampling challenge at this time-point. At day 5, we observed a large expansion of the MuSC and progenitor population (12%) and decline of the myocyte population (1%) as expected. This population heterogeneously expresses lower levels of *Pax7, Vcam1*, and *Sdc4*, higher levels of *Myod1, Des, Mest*, and the mitotic gene *Cdkn1c*, suggesting they include more frequent cells in an activated and/or proliferating state (**Fig. 2C**). This population also induces expression of *Igfbp5*, known to regulate myogenic differentiation (Ren et al., 2008). At day 7, the myogenic cell population remains in a heterogeneously activated and differentiation state with some recovery of *Pax7*^+^ MuSC population (**Fig. 2C**).

Lastly, some cell populations were almost exclusively found at days 0 and 7. We observed a small (3%) population of tenocytes that express the canonical marker *Scx* (scleraxis) as well as *Tnmd* (tenomodulin), *Fmod* (fibromodulin), and *Thbs4* (thrombospondin) (Giordani et al., 2019) (**Fig. S1-3**). We also observed two closely related populations of mesenchymal progenitors and smooth muscle cells, which were distinguished based on their expression of *Myh11* and *Acta2*. In a similar manner, Schwann cells (enriched for *Mpz, Mbp, Fxyd3*, and *Prx*) were distinguished from other related neural and glial cells (enriched for *Ptn, Postn, Cadm1, Lyz2* and *Col20a1*) (**Fig. S1-3**).

### Single-cell trajectory inference organizes a myogenic cell continuum involved in muscle regeneration

The consensus model of adult muscle regeneration states that a subset of MuSCs leave quiescence after tissue damage and enter an activated cycling state to generate progeny through a combination of asymmetric and symmetric division events (Wang and Rudnicki, 2011). These myogenic progenies adopt multiple fate trajectories, including return to *Pax7*^+^ quiescent MuSCs, differentiation into a proliferative *Pax7*^−^ *Myf5*^+^ *Myod1*^+^ myoblast progenitors, or commitment into *Myog*^+^ fusion-competent myocytes. We asked whether unbiased analyses could reconstruct this consensus model and provide into new insights into the continuum of myogenic cell-states.

First, we explored the cellular heterogeneity within the cumulative myogenic cell population by selecting the MuSCs, progenitors, and mature myocytes from the unified transcriptomic atlas. Unbiased SNN clustering revealed five sub-populations of myogenic cells (**Fig. S3A**). These subpopulations clarify the myogenic heterogeneity in two ways: across post-injury time-points and within the stage of differentiation (**Fig. S3B**). Cluster 5 is comprised of terminally differentiated myocytes expressing elevated levels of *Acta1* and *Myh1* from all time-points, whereas clusters 1-4 represent a heterogeneous population of MuSCs and their progeny. Cluster 1 contains cells from both from days 0 and 7 post-injury and is largely enriched for *Pax7, Sdc4, Vcam1*, and *Cd34* transcripts, therefore are likely quiescent MuSCs (**Fig. S3C**). Clusters 2, 3, and 4 contain cells from days 5 and 7 post-injury and that heterogeneously express the myogenic activation and commitment markers *Myf5, Myod1*, and *Myog* and have reduced expression of MuSC markers, suggesting these each contain a mixture of activated MuSC and myoblasts. Notably, clusters 2-4 likely represent a varied but sequential transition towards myogenic commitment, which is exemplified by stepwise elevation in *Myog* expression (**Fig. S3C**). Therefore, we conclude that cluster 4 is comprised of committed myoblasts whereas clusters 2 and 3 primarily contain activated MuSCs. These clusters describe an organized partitioning of the myogenic cell population.

Next, given that SNN clustering lacks hierarchical structure, we sought to organizing these subpopulations using a trajectory inference model to delineate their interrelatedness. We applied Monocle reverse graph embedding (Qiu et al., 2017) to the cumulative myogenic cell population within the atlas to infer a hierarchical trajectory (**Figs. 3A**). The Monocle analysis focused on differentially expressed genes within one of these five myogenic clusters and aligns cells into a one-dimensional “pseudo-time” axis. The pseudo-time trajectory presented an organized, branched progression of cells from quiescent MuSCs to cycling and differentiating progenitors to terminally differentiated myocytes, which can be seen by labelling individual cells using the cell population annotations from the unified atlas (**Fig. 1B**). Both the beginning and ending branches within the trajectory are composed of cells from days 0 and 7 post-injury, indicated they consist of *Pax7*^hi^ quiescent MuSCs and *Acta1*^hi^ terminally differentiated myocytes, respectively, which are both absent at intermediate time-points (**Fig. 3A**). A subset of cells diverts at the central node into a third branch comprised of day 5 and 7 post-injury cells enriched for *Cdk1* and *Cdc20*, indicating that they are actively cycling myoblasts. Near the central node, we observed day 5 and 7 post-injury cells that expressed myoblast and myocyte markers such as *Myog* and *Myod1* (not shown). We interpret this branch structure as a bidirectional trajectory in which activated and cycling progenitors (predominantly from day 5 post-injury) can bifurcate either towards further commitment into terminal myocytes needed for myofiber repair or towards a return to quiescence.

**Figure 3.**
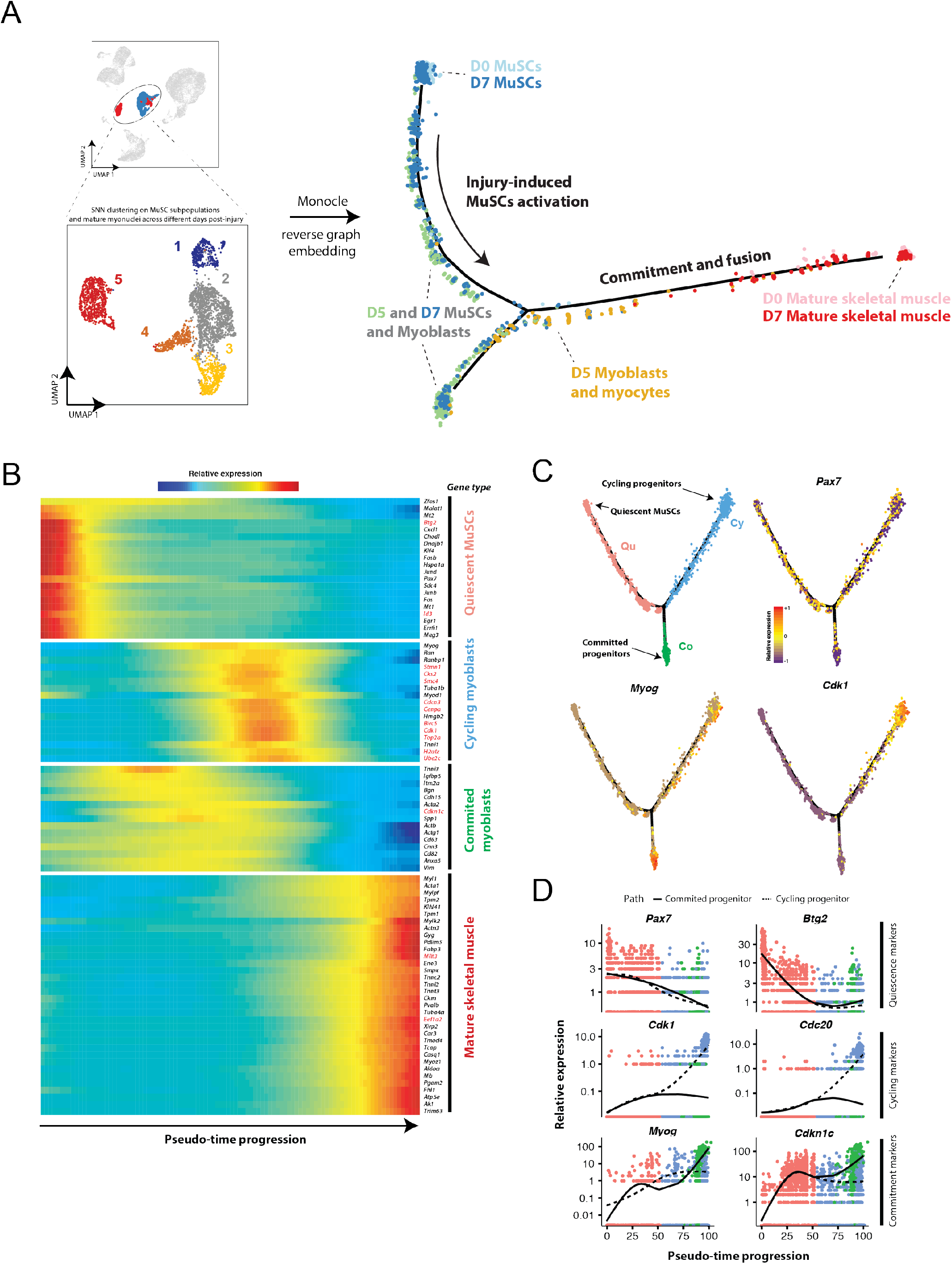
Inferring a muscle stem/progenitor cell hierarchy using Monocle pseudo-time model. (**A**) All cells within the muscle stem/progenitor and mature myocyte cell clusters (3,276 total cells) from day 0, 5, and 7 post-injury (*top left*) were selected and re-analyzed with SNN/UMAP (*bottom left*) and Monocle reverse graph embedding (*right*). Graph embedding results are presented with cells color-coded by day and labeled with cluster identities. (**B**) Heatmap showing four clusters of “pseudo-synchronous” genes that are differentially expressed along the Monocle pseudo-time axis. The color-scale presents the spline-fit relative expression of each gene. Mitotic genes are labeled in red. (**C**) A refined analysis of muscle stem and progenitor sub-populations, after removal of mature myocytes, by Monocle trajectory inference. Monocle feature plots are presented showing three branch groups (*Qu, Cy, Co*) connected by a learned manifold (black lines). Same colors are used to associate individual cells with branch groups in (C, *top left*) and (D). The abundance of *Pax7, Myog*, and *Cdk1* transcripts are plotted for individual cells using a Z-score normalized color-scale. (**D**) Pseudo-time ordered single-cell expression trajectories for genes enriched in the quiescence (Qu) cluster (*Pax7, Btg2*), in the cycling (Cy) cluster (*Cdk1, Cdc20*), and in the commitment (Co) cluster (*Myog, Cdkn1c*). Overlaid lines correspond to inferred cell trajectories associated with ending in the cycling (hatched) and commitment (solid) clusters.

Then, we performed a differential expression analysis along the pseudo-time axis in order to identify genes that explain this myogenic cell progression (**Fig. 3B**). The top 86 differentially expressed genes partition in four clusters that are distinguished by their pseudo-temporal gene expression patterns (see **Methods**). The “early” gene cluster contains quiescence-associated MuSC genes, including *Pax7, Id3* (a direct target of Pax7 and inhibitor of myogenic activation differentiation), *Mt2, Klf4* (a potent inhibitor of smooth muscle differentiation), and cell cycle inhibitor *Btg2* (which has been shown to interact with Id3 to regulate neural progenitor cell differentiation) (Farioli-Vecchioli et al., 2009; Kumar et al., 2009). The first “intermediate” gene cluster group contains activation- and cycling-associated genes, including *Myod1, Hmgb2* (regulates MuSC differentiation through Igf2), as well as multiple mitotic genes such as the cyclin-dependent kinases *Cks2* and *Cdk1, Smc4* (essential to condense chromatin), and the anti-apoptotic gene *Birc5*. The second “intermediate” gene cluster contains commitment-associated genes such as the cyclin-dependent kinase inhibitor *Cdkn1c*, the myogenic differentiation regulator *Igfbp5* (Ren et al., 2008), and genes encoding transmembrane proteins *Cdh15, Itm2a, Cd82* and *Cd63*. The “late” gene cluster contains terminal differentiation-associated genes such as *Myl1*, the troponin family complex members (e.g., *Tnni2*), and metabolic enzymes *Gyg* and *Ak1* (Janssen et al., 2000).

Last, we analyzed the trajectory of the myogenic stem/progenitor cell population (other cell types excluded from this analysis, **Fig. 3C-D** and **S3D**). This refined trajectory model identifies three branches of immature myogenic cells within the transcriptomic atlas (**Fig. 3C-D**), distinguished by their quiescent (“Qu”), cycling (“Cy”), and non-cycling committed (“Co”) signatures. The “Qu” branch is enriched for *Btg2* and *Id3* expression, whereas the “Cy” branch is enriched for G2-state mitotic genes *Cdk1, Cdc20*, and *Ccnb2*, and the “Co” branch, is enriched for the cell cycle inhibitory genes *Cdkn1c* and *Myog* (**Fig. 3D** and **S3E**). When visualized with respect to the pseudo-time progression axis, these genes help distinguish between cells likely to project into “cycling” and “committed” cell fates (**Fig. 3D**). In conclusion, Monocle analysis of the myogenic component of the transcriptomic atlas of regeneration confirms the generally accepted model of adult myogenesis and provides a hierarchy of intermediate myogenic progenitor populations based on cell cycle and commitment gene expression patterns. The clustering and trajectory model together suggest that *Sdc4, Id3*, and *Btg2* are transcriptional markers that aid decomposition MuSC heterogeneity and may regulate distinct MuSC states.

### Diversification of Syndecan receptor expression in myogenic stem and progenitors provides stage-specific heterotypic cell communication channels

Cell communication signals, acting through secreted ligands binding to receptors on muscle stem and progenitor cells, govern a multitude of cell-fate regulation mechanisms critical for muscle homeostasis and regeneration (Yin et al., 2013). To explore the dynamic cell communication network governing muscle repair, we generated a model that scores for interactions between receptors expressed by non-mature myogenic cells and ligands expressed by other cell types. We reasoned that this model could identify ligand-receptor co-expression pairs that provide “insulated” heterotypic cell-cell interactions potentially influencing cell-fate outcomes in myogenic cells but no other cell types in muscle. First, we considered possible ligand-receptor pairs from a database containing 2,009 mouse intercellular interaction signals (Skelly et al., 2018). Second, we identified receptor genes from this list that are differentially expressed in the myogenic stem/progenitor cell populations relative to all other cells within the transcriptomic atlas at any time-point. Third, we calculated interaction scores by multiplying the average transcript expression value of each differentially expressed receptor gene in the myogenic stem/progenitor cell populations by the expression value of each cognate ligand gene (averaged over all cells within each other annotated cell population). We note this model does not consider spatial proximity between cell types, whether proteins are expressed, or whether the putative interaction pairs are documented specifically within myogenic cells.

Our model identifies 63 and 158 ligand-receptor pairs for the uninjured and injured (days 5 and 7 post-injury, combined) muscles, respectively, and 87 of these pairs were unique to injured samples (**Fig. S4**). We only select interactions where the receptor is differentially expressed in the MuSC and progenitor population. Moreover, for each pair, we consider the interaction significant when the score is greater than the 50^th^ percentile when compared to all cell types. We represent significant interactions by a pairwise chord plot (**Fig. 4A**). In uninjured muscles, the majority of these pairs involve myogenic cell expression of receptor genes *Fgfr1, Fgfr4, Sdc4, Tgfbr3, Cd63* or *Cd82*, consistent with findings that MuSCs express diverse members of the FGFR, Syndecan (Sdc), TGF-β and tetraspanin families (Pawlikowski et al., 2017). Notably, *Fgfr1* and *Fgfr4* interactions are mediated by a diverse set of 15 FGF ligand genes expressed across multiple cell types (mature skeletal myocytes, FAPs, tenocytes, and neural/glial/Schwann cells), suggesting broad redundancy in FGFR signaling interactions in uninjured muscles (**Fig. S4**). This ligand-receptor promiscuity is also exemplified by the pairs involving *Sdc4*, the only *Sdc* gene differentially expressed by myogenic cells with uninjured muscles, which involve multiple ligand genes from various cell types including *Ccl5* (myeloid progenitors), *Thbs1* and *Tnc* (tenocytes), *Fgf2, Tfpi, Mdk*, and *Rspo3* (FAPs), *Tgm2* and *Cxcl12* (SMCs), and *Adam12* (neural/glial cells), and *Fgf6* (mature myocytes). In the injured muscle we observed a diversification in the ligand-ligand interaction landscape, highlighted by inclusion of additional myogenic cell receptor genes (*Sdc1, Sdc2, Notch1, Notch3*, and *Cd151*) within the interaction pairs.

**Figure 4.**
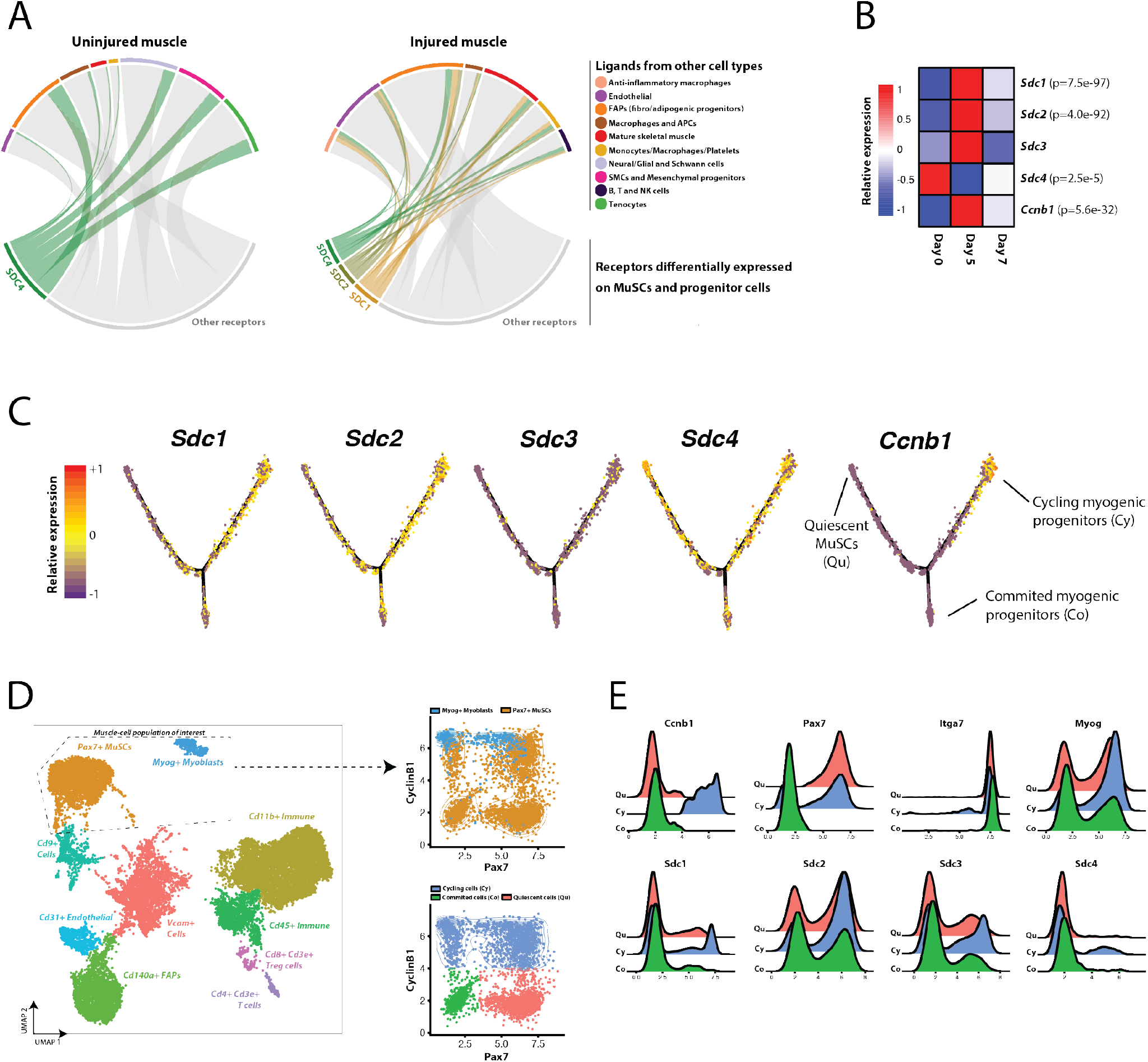
Ligand-receptor model reveals diversification of communication signals through Syndecan family receptors during muscle regeneration. (**A**) Chord plot summarizing the significant pairwise interactions between receptor genes that are differentially expressed in the myogenic stem/progenitor population and ligand genes expressed by other cell types within the transcriptomic atlas. Left, uninjured (day 0) samples. Right, injured (day 5 and 7 post-injury) samples. Differentially expressed receptor genes outside of the Syndecan family are in grey. For a given receptor-ligand pair, we only represent interactions whose score (Fig. S4) is greater than the 50^th^ percentile across all cell types. (**B**) *Sdc1/2/3/4* and *Ccnb1* (Cyclin-B1) transcript averages across all non-mature myogenic cells within the transcriptomic atlas, split by days post-injury (see **Fig. 3C**). *p*-values listed if differentially expressed across time-points when modeled using a negative binomial distribution. (**D**, left) CyTOF atlas, consisting of 19,028 cells collected from regenerating (day 5 post-injury) muscles and stained with a panel of 35 antibodies (see **Table S1**) including Syndecan-1/2/3/4 and Cyclin-B1. UMAP and unsupervised SNN clustering identified 11 populations including a population of Pax7^+^ MuSCs (orange) and Myog^+^ myogenic progenitors (blue). These two myogenic clusters were grouped for further analysis. (**D**, right) Cyclin-B1 versus Pax7 scatter plots. Top coded using CyTOF SNN cluster identifiers. Bottom, coded by sub-population gates: Cyclin-B1^−^ Pax7^+^ quiescent cells (Qu; pink), Cyclin-B1^+^ cycling progenitors (Cy; blue), and Cyclin-B1^−^ Pax7^−^ committed myocytes (Co; green). (E) expression histograms for Syndecan-1/2/3/4 and other myogenic markers for the three subpopulations identified in (D).

Given the observed frequent and diverse involvement of *Sdc* receptor genes in these coexpressed ligand-receptor scores and their documented role in MuSC regulation (Pisconti et al., 2012), we asked whether *Sdc* genes exhibit stage-specific expression patterns within the myogenic compartment of the transcriptomic atlas. We performed differential expression testing on the non-mature myogenic cell populations within the transcriptomic atlas between the day 0, 5, and 7 samples, and found that *Sdc1* and *Sdc2* were elevated at day 5 post-injury (along with the cycling MuSC gene *Ccnb1*), and *Sdc4* was elevated in the uninjured muscle (**Fig. 4B**). Further, *Sdc1* and *Sdc2* have enhanced expression in the *Ccnb1*^hi^ “Cy” branch of the Monocle trajectory (**Fig. 4C**).

We validated Syndecan expression variation at different stages of the myogenic hierarchy through an independent analysis based on multiparameter mass cytometry (CyTOF). We immunostained a single-cell suspension from digested muscles at 5 days post-notexin injury using a panel of 35 antibodies (including the four Sdcs) to label myogenic and other cell types. After gating for live cells and removing debris (**Fig. S5**), the CyTOF data was analyzed using a similar bioinformatic pipeline as for scRNA-seq datasets. Unsupervised SNN clustering revealed 11 subpopulations of cells including endothelial cells, FAPs, macrophages, lymphocytes, mesenchymal progenitors, an unresolved group of Cd9^+^ cells, and two myogenic populations (**Fig. 4D**). One myogenic population consists of Pax7^+^ MuSCs whereas the other consists of Myog^+^ myoblasts and myocytes (**Fig. 4D**). These myogenic populations were merged, and sub-populations were gated into Cyclin-B1^−^ Pax7^+^ quiescent cells, Cyclin-B1^+^ cycling progenitors, and Cyclin-B1^−^ Pax7^−^ committed myocytes (**Fig. 4E**). We observed that these three sub-populations uniformly express the myogenic surface marker Integrin-α7 but have heterogeneous expression of Sdcs. In particular, Sdc-1^hi^ and -2^hi^ cells are enriched in the cycling progenitor sub-populations. Taken together, these observations suggest that the transient induction of Sdc-1 and -2 in cycling (Cyclin-B1^+^) muscle progenitors may allow for diversified engagement with an exclusive set heterotypically secreted ligands in regenerating, but not uninjured, muscles.

## Discussion

Combined with the development of increasingly complex computational methods, scRNA-seq has emerged as a powerful tool to profile the transcriptome of thousands of individual cells in one experiment (Stuart and Satija, 2019). scRNA-seq analysis permits an unbiased survey of cellular complexity and heterogeneity with substantial experimental scope. Here, we leverage these recent developments to build a comprehensive temporal atlas of muscle tissue repair with over 34,000 single-cell transcriptomes, adding to the growing repository of single-cell skeletal muscle datasets.

To date, a handful muscle-focused scRNA-seq projects have been reported. The first study was performed on FACS-sorted *Pax7*-tdTomato^+^ MuSCs using the Fluidigm C1 system (Cho and Doles, 2017). Though limited to 21 single MuSCs, it provided a transcriptomic view of MuSC heterogeneity, highlighted enriched levels of *Cd34* and *Sdc4* transcripts. The study also highlighted some technical limitations, low recovery of *Pax7* transcripts, which motivates the need to generate greater numbers of single-cell transcriptomes to allow more robust statistical analyses. Recently, Giordani et al. presented a transcriptomic atlas of 12,441 muscle-resident cells (Giordani et al., 2019). Their study identified 10 distinct types of cells, including a population of Itga7^+^ Vcam1^−^ smooth muscle and mesenchymal cells that enhance MuSC engraftment when co-transplanted in mice. Their findings illustrated the potential of high-throughput single-cell analysis to reveal poorly described populations and to generate new hypotheses. Whereas the Giordani et al. study focused on homeostatic mouse muscle tissue, another recent scRNA-seq study focused on regenerating muscle. Dell’Orso et al. presented a transcriptomic atlas of about 3,500 FACS-sorted MuSCs and progenitor cells from homeostatic and notexin-injured muscles (Dell’Orso et al., 2019). They identified two subpopulations of MuSCs with distinct yet overlapping gene expression profiles corresponding to a quiescent and activated state. In addition, they aligned in pseudo-time injured, uninjured MuSCs, and primary myoblasts to reveal 7 classes of genes, including of mitochondrial and glycolytic origin, from which they inferred dynamics in metabolic reprogramming activity. Though the first report to describe the dynamics of MuSCs activation and differentiation by scRNA-seq, their analysis was limited to FACS-sorted cells and a single time-point postinjury (60 hours), which might omit the cellular and temporal complexity of muscle regeneration. Finally, using a CyTOF dataset composed of 23 markers, Porpiglia et al. built a trajectory model using the X-shift algorithm of myogenic differentiation post-injury (Porpiglia et al., 2017). They identified two new surface markers, CD9 and CD104, that were used to describe two subpopulations of muscle progenitors, demonstrating how trajectory models from single-cell data can be used to discover new combinations of surface markers for the prospective isolation of MuSCs and their progeny.

Here, we present a unified and annotated single-cell transcriptomic reference atlas of muscle regeneration in adult mice. Our scRNA-seq and CyTOF analyses confirm prior consensus regarding the cell populations involved in the temporal response to muscle injury, and providing a deeper annotation of additional cell types, sub-populations and states with more resolved dynamics, compared to prior scRNA-seq studies. We present comprehensive scRNA-seq dataset compendium describing a total of 34,438 cells (15 different cell types via SNN clustering) including 3,276 from MuSCs and mature muscle cells (**Figs. 1** and **2**). The complexity of this transcriptomic atlas powered development of a hierarchical continuum model of myogenic cell populations and receptor-ligand cell communication analysis (**Figs. 3** and **4**). We also presented a CyTOF dataset composed of 35 markers to provide an orthogonal validation of myogenic sub-populations and their surface receptor expression variability. Trajectory analysis allowed us to parse the myogenic differentiation lineage post-injury in four distinct groups: quiescent MuSCs, cycling progenitors, committed progenitors, and mature skeletal muscle, with distinct gene expression signatures (**Fig. 3**). Here we resolved that *Id3* and *Btg2* are both enriched in quiescent MuSCs with their expression decreasing following injury-induced MuSC activation and differentiation. Id3 is a DNA binding protein that has been found to be a direct target of Pax7 (Kumar et al., 2009). Id3 is robustly expressed in quiescent MuSCs and blocks differentiation either by directly blocking the activity of pro-myogenic transcription factors such as Myf5, MyoD or by maintaining high levels of Hes1 (Kumar et al., 2009). The role of Btg2 has not yet been previously described in myogenic cells. In hematopoietic and neural lineages, Btg2 promotes differentiation by inhibiting both Id3 and cyclin D1 to restrict cell cycle progression (Yuniati et al., 2019). Though we did not identify the mechanism of Btg2 regulation of myogenic differentiation, these data suggest *Btg2* and *Id3* are distinct transcriptional markers of quiescent MuSCs.

We developed a cell communication model that allowed us to map interactions between MuSC receptors and ligands expressed by other cell types during muscle repair, which highlighted the complex role of Syndecan receptors in coordinating muscle progenitor heterogeneity. Syndecans (Sdcs) are transmembrane heparan sulfate proteoglycans that have been characterized as regulators of muscle development, homeostasis, and regeneration (Pisconti et al., 2012). Previous studies have found that *Sdc1* is only expressed in developing muscle, while the other muscle Sdcs (*Sdc2, Sdc3, Sdc4*) are expressed in MuSCs with differing post-injury expression dynamics specific to each receptor (Pisconti et al., 2012). *Sdc3* plays a role in maintaining MuSC quiescence, as *Sdc3^-/-^* muscles have do not atrophy, despite showing a loss in MuSC number, but have a homeostasis defect (Pisconti et al., 2016; Pisconti et al., 2010). *Sdc4*, in contrast, seems to play a role in MuSCs activation through regulating FGF and HGF signaling (Cornelison et al., 2004). Both our scRNA-seq and CyTOF data suggest that these four Sdcs are expressed heterogeneously with quiescent, cycling, and committed myogenic stem/progenitor cells (**Fig. 4**). *Sdc1*, though previously thought not to be expressed in postnatal muscle, was detected both at the transcription and protein level, by scRNA-seq and CyTOF respectively, in activated cycling muscle progenitors and committed cells but not in MuSCs. Moreover, *Sdc2* was expressed in some quiescent MuSCs and activated progenitors, *Sdc3* expression was restricted to cycling MuSCs progenitors, and *Sdc4* expressed in quiescent MuSCs and cycling progenitors alike. Together, these Sdcs provide a significant fraction of the heterotypic cell communication interactome, suggesting that temporal heterogeneity in Sdc expression may enable myogenic stage-specific fate regulation to a shared set of Sdc-binding ligands.

Though clustering and visualization approaches presented here and by others can be used to identify new cell types and biomarkers, we urge caution on how cell types are defined from these data. First, single-cell data is not immune to technical imperfections such as doublets or sequencing reads that misalign (Stuart and Satija, 2019). Second, single-cell data is sensitive to sample preparation methods, especially from the solid tissue sources. For example, we observed significant variability in the inclusion of tenocytes based on muscle-tendon resection precision (data not shown). We also found other biases introduced in the scRNA-seq composition introduced by FACS sorting can select for metabolically active immune cells. Furthermore, since scRNA-seq is based on RNA detection, some quiescent cell types such as MuSCs are challenging to unambiguously detect and will be benefited from *in vivo* transcript recovery advances (Machado et al., 2017; van Velthoven et al., 2017). Even with these limitations, our scRNA-seq study provides a view that discards some prior assumptions on the boundaries between myogenic cell types and suggests a myogenic continuum that is endowed with a plasticity of cell-cycle and commitment states. This annotated temporal scRNA-seq atlas of muscle regeneration may provide reference resource to examine the role cellular diversity and communication in aging, disease, and across species.

## Methods

### Mice and single-cell isolation

The Cornell University Institutional Animal Care and Use Committee (IACUC) approved all animal protocols and experiments were performed in compliance with its institutional guidelines. Adult C57BL/6J mice were obtained from Jackson Laboratories (#000664, Bar Harbor, ME) and were used at 4-5 months of age. To induce muscle injury, both tibialis anterior muscles received a single 10 μL intramuscular injection of notexin (10 μg/ml; Latoxan, France). At either 2, 5, or 7 days post-injury, one tibialis anterior muscle per mouse was collected to generate single-cell suspensions. Muscle were digested with 2.5 mg/ml Collagenase D (Roche) and 0.04 U/ml Dispase II (Roche) followed by manual dissociation, filtration, and red blood cell lysis to generate a single-cell suspension, as previously described (Cosgrove et al., 2014; Sacco et al., 2008).

### FACS sorting

We used a fluorescence-activated cell sorter (FACS ARIA Fusion, BD Biosciences) to enrich the single-cell suspension for Calcein-AM^+^ propidium iodide^−^ viable cells. The single-cell suspension was stained with Calcein-AM (1 μg per 100,000 cells) and PI for 30 min in PBS. Live cells were sorted selecting for Calcein-AM^+^ and debris were removed using a standard FSC/SSC gating strategy.

### Single-cell RNA-sequencing

After digestion, single-cells were washed and resuspended in 0.04% BSA in PBS at a concentration of at least 400 cells/μL. Cells were counted manually with a hemocytometer in order to determine the concentration. The single-cell RNA-sequencing library was then prepared using the Chromium Single Cell 3’ reagent kit v2 (10X Genomics) in accordance with the manufacturer’s protocol. Briefly, the cells were diluted into the Chromium Single Cell A Chip as to yield a recovery of ~6,000 single-cell transcriptomes with <5% doublet rate. Following the library preparation, the libraries were sequenced in multiplex (n=2 per sequencing run) on the NextSeq 500 sequencer (Illumina) to produce between 200 and 250 million reads per library and on average a minimum of 30,000 reads per single-cell.

### Analysis of scRNA-seq data

Sequencing reads were processed with the Cell Ranger version 3.0.1 (10X Genomics) using the mouse reference transcriptome mm10. From the gene expression matrix, the downstream analysis was carried out with R version 3.5.1 (2018-07-02). Quality control, filtering, data clustering and visualization, and the differential expression analysis was carried out using Seurat version 2.3.4 R package (Butler et al., 2018) with some custom modifications to the standard pipeline. Each of the 9 datasets were first analyzed independently before combining datasets from the same timepoint together for an integrated analysis. For each individual dataset, genes expressed in less that 3 cells as well as cells <1000 UMIs and <200 genes were removed from the gene expression matrix. In addition, we removed any single-cell with >20% UMIs mapped to mitochondrial genes, as well as obvious outliers in number of UMIs (cell doublets). After log-normalizing the data, the expression of each gene was scaled regressing out the number of UMI and the percent mitochondrial gene expressed in each cell. We performed PCA on the gene expression matrix and used the first 15 principal components for clustering and visualization. Unsupervised shared nearest neighbor (SSN) clustering was performed with a resolution of 0.4 and visualization was done using uniform manifold approximation and projection (UMAP) (Becht et al., 2018). Finally, differential expression analysis was achieved using Seurat’s “FindAllMarkers” function using a likelihood ratio test that assumes the data follows a negative binomial distribution and only considering genes with >log_2_(0.25) fold-change and expressed in at least 25% of cells in the cluster.

To analyze datasets merged from the same timepoint, we performed canonical correlation analysis (CCA) between datasets followed by data alignment using nonlinear dynamic time warping. We used the CCA matrix instead of PCA for visualization and clustering, and consequently, the differential expression analysis between clustered cells. The CCA space allows us to highlight shared patterns in gene expression profile between datasets. In addition, we did not observe significant differences in results between working in the CCA or PCA space, thus indicating little batch effect between samples from the same timepoint. We finally combined the 9 datasets into a uniform UMAP atlas using the CCA scores calculated from all the datasets. However, we retain the labels identified independently at each timepoint and the data was not re-clustered.

### Monocle trajectory analysis

We used the Monocle version 2.8.0 R package (Qiu et al., 2017) to organize cells in pseudo-time and infer new trajectories of MuSCs subpopulations post-injury. First, we subsetted the cells labeled as MuSCs and progenitor cells and Mature skeletal muscle from the Seurat dataset and across all timepoints and samples. Second, we performed unsupervised SNN clustering in order to identify new subpopulations in the data, from which we then used the Seurat “FindAllClusters” function (as described above) to find differentially expressed genes that characterize the subpopulations. We then selected the top 150 genes based on fold-change expression with a minimum of log_2_(0.8) and adjusted p-value of 0.1. This list genes of differentially expressed genes is then used by Monocle for clustering and ordering cells using the DDRTree method and reverse graph embedding. To identify genes that are differentially expressed across Monocle branches (States), we transferred the labels back to the Seurat dataset and performed differential expression analysis as described above.

### Ligand-receptor cell communication model

The model aims at scoring potential receptor-ligand interactions between MuSCs (receptor) and other cell types (ligand). We used the receptor-ligand interaction database from Skelly et al. (Skelly et al., 2018). To calculate the score for a given receptor-ligand pair, we multiply the average receptor expression in MuSCs with the average ligand expression per other cell type (including MuSCs to consider for autocrine interactions). We only considered receptors that are differentially expressed in MuSCs at any given time-point post-injury.

### CyTOF sample preparation and staining

Most of the cell surface markers were obtained from Fluidigm and few others were conjugated in-house using the Maxpar X8 Multimetal Labeling Kit (see **Table S1** for complete antibody list). The cells were stained with 50 μl of the antibody cocktail (100 μl of total staining volume) for 30 minutes at RT with intermittent vortexing. Following 2 washes (300g, 5 mins, RT), the cells were fixed with freshly prepared 1.6% PFA for 10 minutes. Thereafter, the cells were incubated for 30 minutes in 1 mL of the Nuclear Antigen Staining Buffer working solution. Then the cells were washed with 2 ml of Nuclear Antigen Staining Perm at 800g, 5 minutes, RT. 50 μl of nuclear antigen antibody cocktail was added to 50 μl of cell pellet solution and incubated for 45 minutes at RT. Following antibody staining, the cells were also stained with Cell-ID Intercalator-Ir-125 μM diluted to 1:1000 with MaxPar Fix and Perm buffer for 1 hour at RT. In addition, cells were also stained with 10 μl of Cisplatin for viability in 1 mL of pre-warmed serum free medium for 5 minutes at RT. Finally, cells were also stained with IdU (5-Iodo-2-deoxyuridine) to label the S-phase at a concentration of 50 μM for 30 minutes at 37°C.

### CyTOF data acquisition and analysis

Cells were washed twice with staining buffer and then with MilliQ water. After the final wash, the cells were adjusted to a concentration of 10^6^ cells/mL with 1:10 EQ beads to MilliQ water solution. Prior to the acquisition, the instrument was tuned and calibrated using the EQ standard beads. The acquisition speed of the sample was maintained within 400 events/seconds to avoid doublets and ion cloud fusion errors in the data. The output FCS files were normalized using the Fluidigm normalizer algorithm that is embedded within the CyTOF software (Version 6.7.1014). The CyTOF data was first gated using the Cytobank software in order to exclude debris, dead cells and doublets (Kotecha et al., 2010). The resulting FCS file was then converted into a gene expression matrix using the Cytofkit R package using an inverse hyperbolic sine transformation (Chen et al., 2016). The gene expression matrix was then analyzed with a Seurat-based custom pipeline, which allowed for SSN clustering and UMAP visualization.

### Reagents and Resources

| **Antibodies** |  |  |
| --- | --- | --- |
| Anti-Mouse CD8a (53-6.7) | Fluidigm | 3153012B |
| Anti-Mouse CD9 (KMC8) | Fluidigm | 3158009B |
| Anti-Mouse CD11b (M1/70) | Fluidigm | 3143015B |
| Anti-Mouse CD3e (145-2C11) | Fluidigm | 3152004B |
| Anti-Mouse CD4 (RM4-5) | Fluidigm | 3145002B |
| Anti-Mouse CD45.2 (104) | Fluidigm | 3147004B |
| Anti-Mouse CD206/MMR (C068C2) | Fluidigm | 3169021B |
| Anti-Mouse CD80 (16-10A1) | Fluidigm | 3171008B |
| Anti-Mouse CD31/PECAM-1 (390) | Fluidigm | 3165013B |
| Anti-Mouse CD140a (APA5) | Fluidigm | 3148018B |
| Anti-Mouse CD140b (APB5) | Fluidigm | 3151017B |
| Anti-CyclinB1 (GNS-1) | Fluidigm | 3164010A |
| Cell-ID™ IdU | Fluidigm | 201127 |
| Anti-Mouse CD15 (mc-480) | Lederer Lab -- Harvard Medical School | http://ledererlab.bwh.harvard.edu/cytof-core/ |
| Anti-Mouse CD29 (HMB1-1) | Lederer Lab -- Harvard Medical School | http://ledererlab.bwh.harvard.edu/cytof-core/ |
| Anti-Mouse CD34 (MEC14.7) | Lederer Lab -- Harvard Medical School | http://ledererlab.bwh.harvard.edu/cytof-core/ |
| Anti-Mouse CD106/VCAM-1 (429) | Lederer Lab -- Harvard Medical School | http://ledererlab.bwh.harvard.edu/cytof-core/ |
| Anti-Mouse Integrin alpha 7 | R&D Systems | MAB3518 |
| Anti-Human/Mouse/Rat/Chicken Pax7 | R&D Systems | MAB1675 |
| Anti-Human/Mouse Myogenin | R&D Systems | MAB6686 |
| Anti-Human/Mouse Myosin Heavy Chain Antibody | R&D Systems | MAB4470 |
| Anti-Mouse VE-Cadherin | R&D Systems | MAB1002 |
| Anti-Mouse Laminin alpha 1 | R&D Systems | MAB4656 |
| Anti-Mouse Sca-1/Ly6 | R&D Systems | AF1226 |
| Anti-Mouse C1q R1/CD93 | R&D Systems | MAB1696 |
| Anti-Human/Mouse CX3CR1 | R&D Systems | AF5825 |
| Anti-Mouse Mouse Syndecan-2/CD362 | R&D Systems | AF6585 |
| Anti-Human/Mouse MyoD (5.8A) | Novus Biologicals | NB100-56511 |
| Anti-Human/Mouse/Rat CD82 | Thermo Fisher Scientific | PA5-20356 |
| Anti-Mouse COL1A1 (3G3) | Santa Cruz Biotechnology | sc-293182 |
| Anti-Mouse Syndecan-3 Antibody (G-2) | Santa Cruz Biotechnology | sc-398194 |
| Anti-Mouse CD138/Sdc1 (281-2) | BD | 553712 |
| Anti-Mouse Syndecan-4 (KY/8.2) | BD | 550350 |
| Anti-Mouse Scx (N-term) | Abgent | AW5469-U100 |
| Anti-Mouse Fibromodulin (3E9D10) | Proteintech | 60108-1-Ig |

| **Reagents** |  |  |
| --- | --- | --- |
| Cell-ID Intercalator-Ir | Fluidigm | 201192A |
| Cell-ID IdU | Fluidigm | 201127 |
| Cell-ID Cisplatin | Fluidigm | 201064 |
| Maxpar Cell Staining Buffer | Fluidigm | 201068 |
| Dispase II (neutral protease, grade II) | Sigma-Aldrich | 04942078001 |
| Collagenase D, from Clostridium histolyticum | Sigma-Aldrich | 11088866001 |

| **Commercial kits** |  |  |
| --- | --- | --- |
| Chromium Single Cell 3' Library & Gel Bead Kit v2 | 10X Genomics | CG00052 (protocol) |
| Maxpar X8 Multimetall Labeling Kit | Fluidigm | 201300 |

| **Deposited data** |  |  |
| --- | --- | --- |
| Mouse ligand-receptor database | Skelly et al., 2018 | https://www.ncbi.nlm.nih.gov/pubmed/29346760 |
| scRNA-seq data | This paper |  |
| CyTOF data | This paper |  |

| **Software packages and algorithms** |  |  |
| --- | --- | --- |
| Cytofkit | Chen et al., 2016 | https://github.com/JinmiaoChenLab/cytofkit |
| Cell Ranger 3.0 | 10X Genomics | https://support.10xgenomics.com/single-cell-gene-expression/software/downloads/latest |
| CyTOF Software 6.7 | Fluidigm | https://www.fluidigm.com/software |
| Cytobank | Cytobank, Inc. | https://www.cytobank.org |
| Seurat 2.3.4 | Butler et al., 2018 | https://github.com/satijalab/seurat |
| Monocle 2.8.0 | Qiu et al., 2017 | https://github.com/cole-trapnell-lab/monocle-release |

## Acknowledgments

This work was financially supported by National Institutes of Health under awards R00AG042491 (to B.D.C), R01AG058630 (to B.D.C.), and 1DP2AI138242 (to I.D.), a US Department of Education Graduate Assistantship in Areas of National Need under Award P200A150273 (to A.J.D.), a Glenn Medical Research Foundation and American Federation for Aging Research Grant for Junior Faculty (to B.D.C.), and Cornell Engineering Learning Initiatives Undergraduate Research Award (to P.F.). The authors acknowledge technical assistance from Peter Schweitzer of Genomics Facility at the Cornell University Biotechnology Resource Center. The authors are also thankful for the constructive discussions with members of the Cosgrove, De Vlaminck, and Elemento research groups, as well as Christopher Mendias, Laura Donlin, and their research groups at the Hospital for Special Surgery.

## Author Contributions

A.J.D. and B.D.C. designed the study and wrote the manuscript. A.J.D., P.F., and S.S.B. organized the mouse colony and procedures and performed the scRNA-seq study. A.J.D. and H.R. performed the CyTOF study. A.J.D. performed the scRNA-seq and CyTOF data analysis, with supervision and assistance from B.D.C., I.D., and O.E. All authors reviewed and edited the manuscript.

## Supplementary Material

**Figure S1-1.**
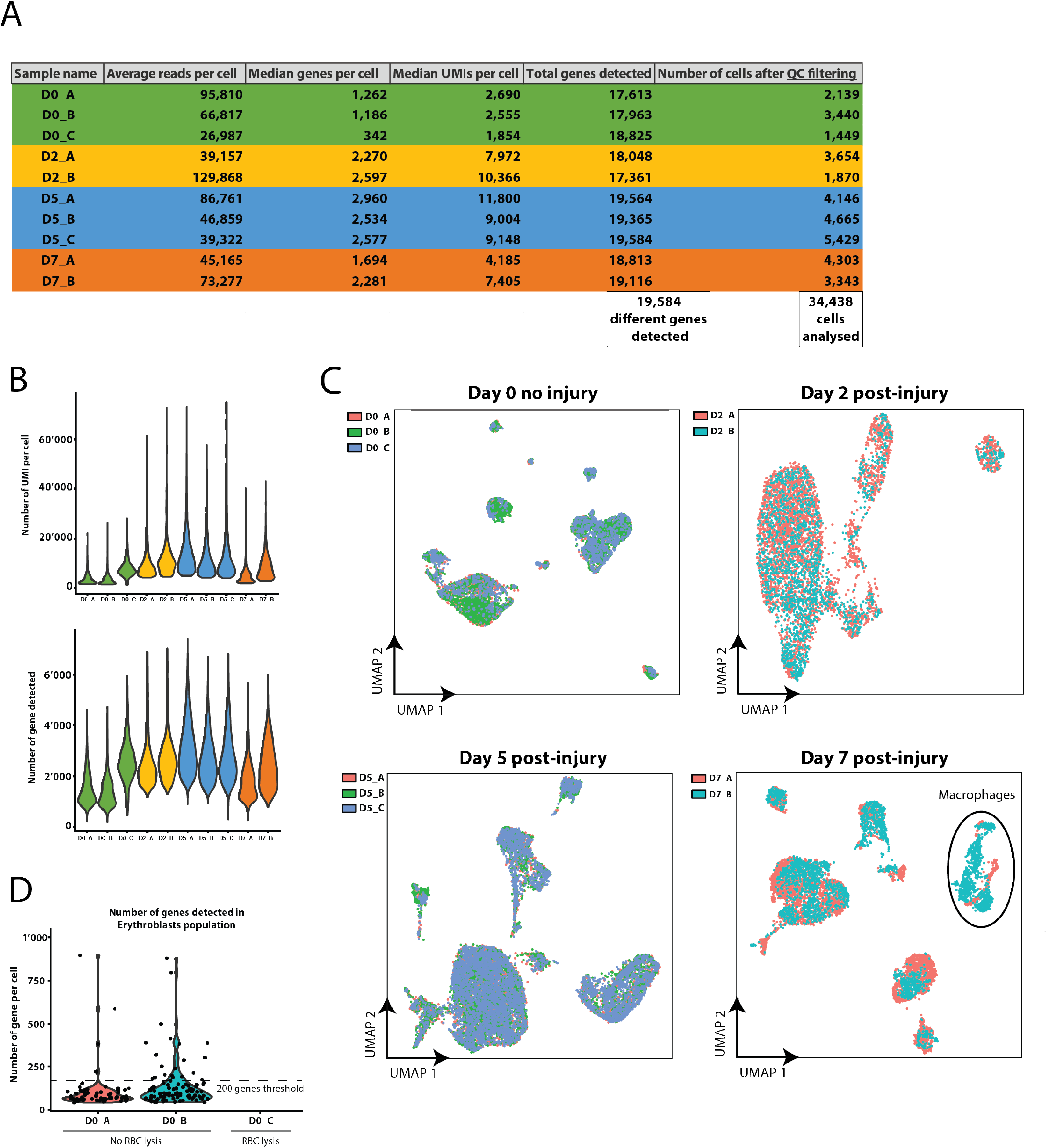
Technical and quality control measures for scRNA-seq datasets. (**A**) Sequencing reads processing statistics from the 10X Cell Ranger pipeline. The samples originated from 9 individual female mice at 4-6 months of age. Samples are color-coded by time-point. Differences in sample quality were addressed by applying a quality control (QC) filter in Seurat. The last column represents the number of cells that were used for the analysis after QC. A total number of 34,438 cells expressing 19,584 unique genes were used for the downstream analysis. (**B**) Number of unique molecular identifiers (UMIs) and genes per cell per sample after QC filtering. On average cells have more UMIs and genes detected for injured samples (day 2 and 5 post-injury). The samples are color-coded as in (A). (**C**) Technical effect and differences in population number across samples from the same timepoint. Single-cell transcriptomes were projected on a UMAP plot using cross-correlation analysis (CCA) scores between samples. Some divergences in cell number can be observed for the immune cluster between D7_A and D7_B. (**D**) Distribution of the number of genes per Erythroblast cell type with or without red blood cell (RBC) lysis during the single-cell suspension preparation. Erythroblasts express a small number of genes and were removed from the analysis after applying a 200-gene minimum filter. RBC lysis also removed all detectable Erythroblasts. RBC lysis was applied to all samples in this study expect D0_A and D0_B.

**Figure S1-2.**
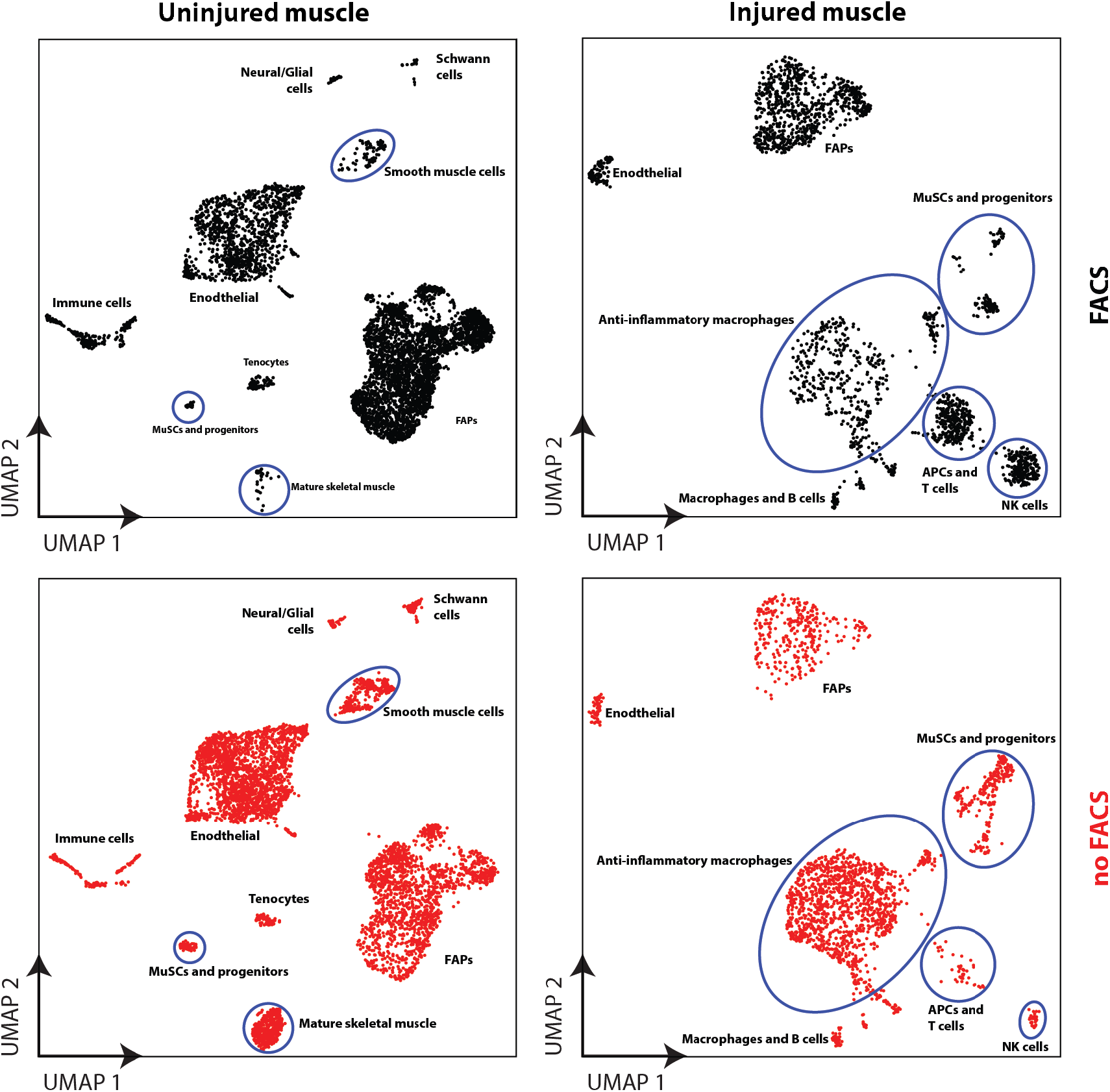
Evaluation of selection biases and resulting cell type abundances introduced by FACS filtering prior to scRNA-seq. UMAP projection of single-cell transcriptomes obtained by either the standard cell isolation protocol (see **Methods**) or to one that included a Calcein-AM^+^ enrichment of live cells and removal of debris by FACS. FACS selection introduces measurable biases in cell population number for smooth muscle cells, MuSCs, and mature skeletal muscle in the uninjured muscle and various immune cell types in the injured muscle.

**Figure S1-3.**
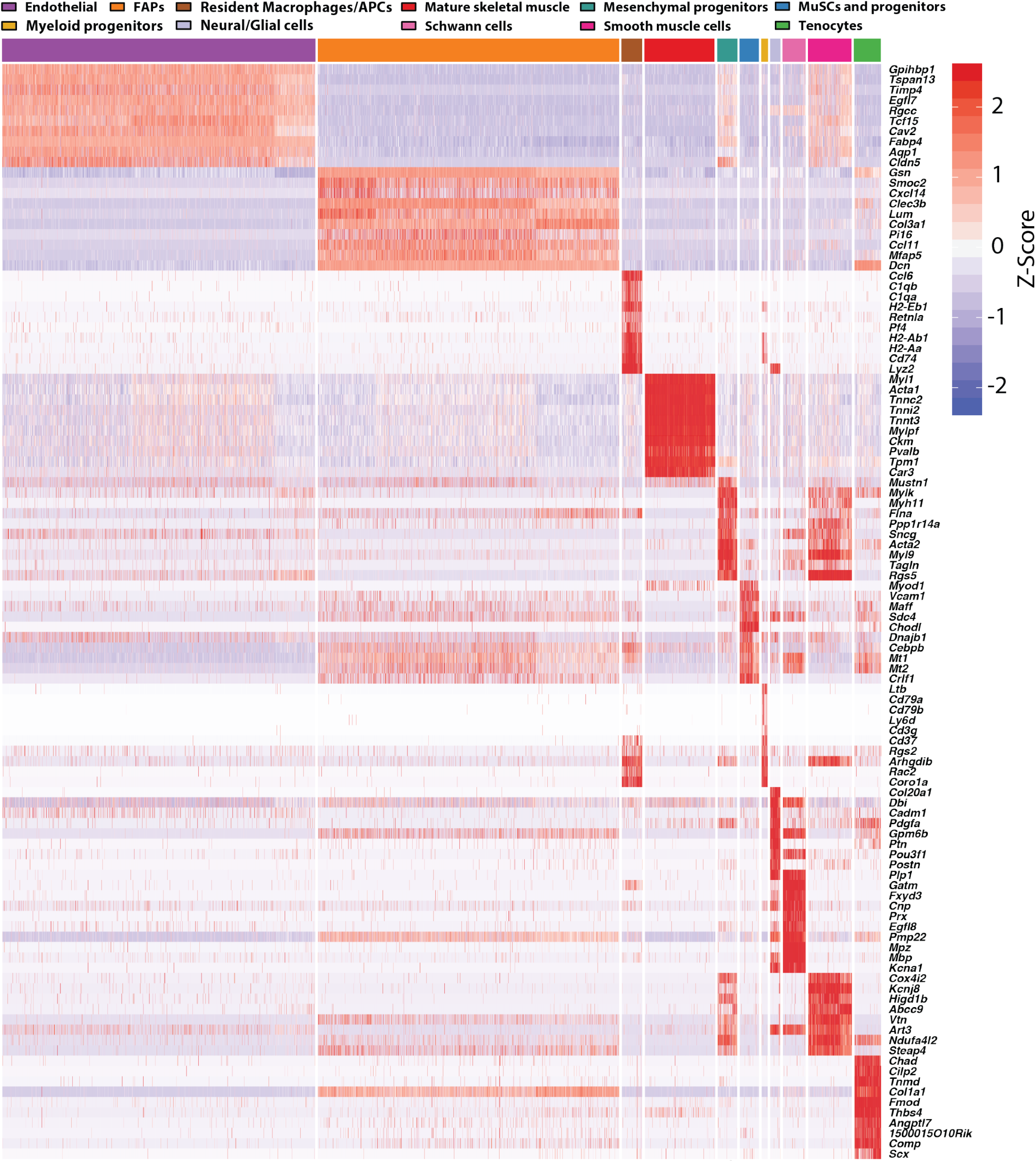
Top differentially expressed genes per cluster in the uninjured (day 0) muscle. Normalized expression (Z-score) heatmap for top differentially expressed genes in the 11 subpopulations identified by SNN at the uninjured (day 0) time-point. The columns represent cells and are organized by cell type as color-coded.

**Figure S1-4.**
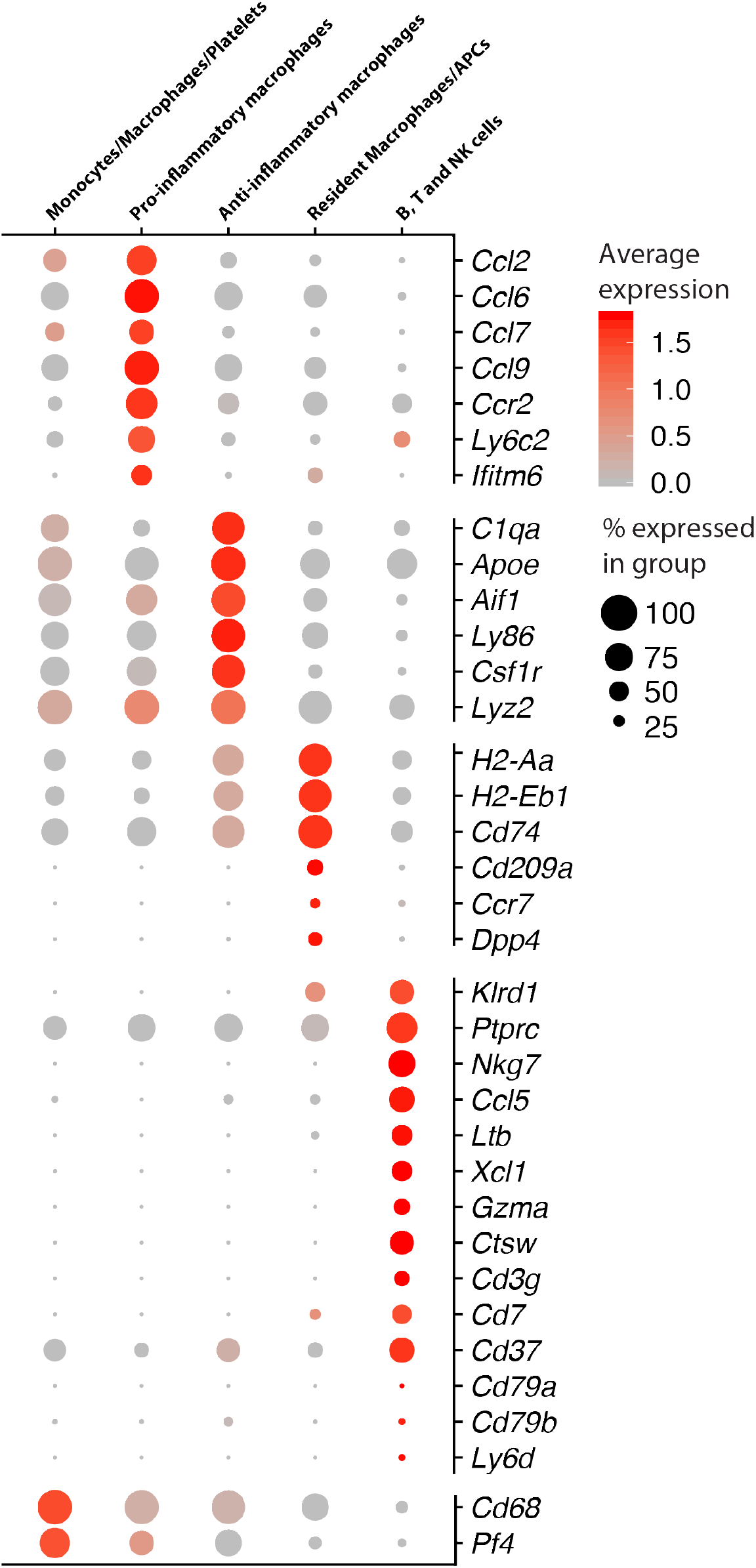
Expression of immune markers identified by differential expression analysis. Average expression of immune genes identified by differential expression analysis in the combined dataset (**Fig. 1B**). These markers allowed delineation of 5 different immune cell populations pre and post-injury. The dot size represents the percentage of cells within a group with an expression level >0.

**Figure S2-1.**
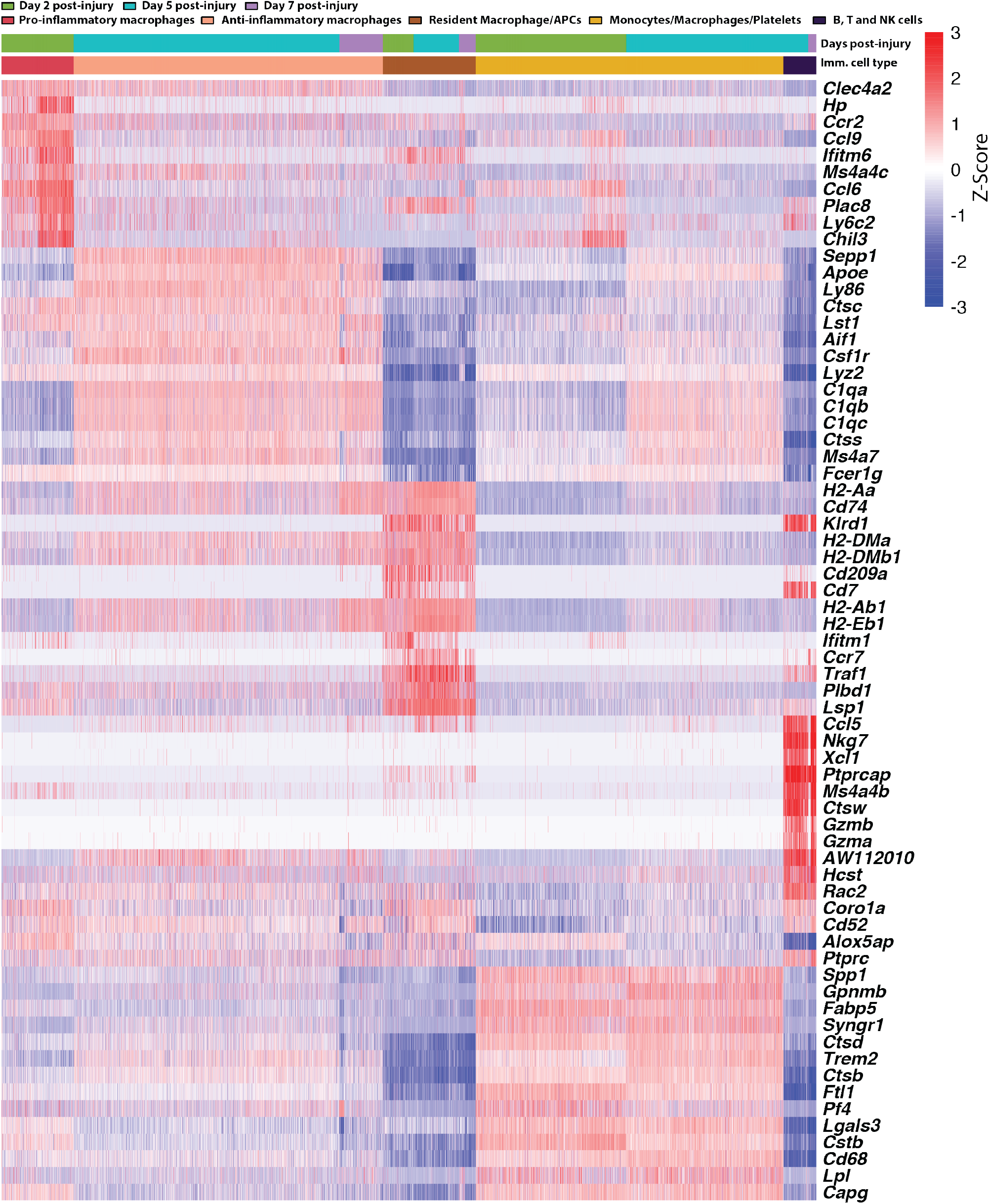
Top differentially expressed genes in immune subpopulations across days post-injury. Normalized expression (Z-Score) heatmap for top differentially expressed genes in 5 subpopulations of immune cells identified by SNN post-injury. The columns represent cells and are organized first by cell type and then by day post-injury as color-coded.

**Figure S2-2.**
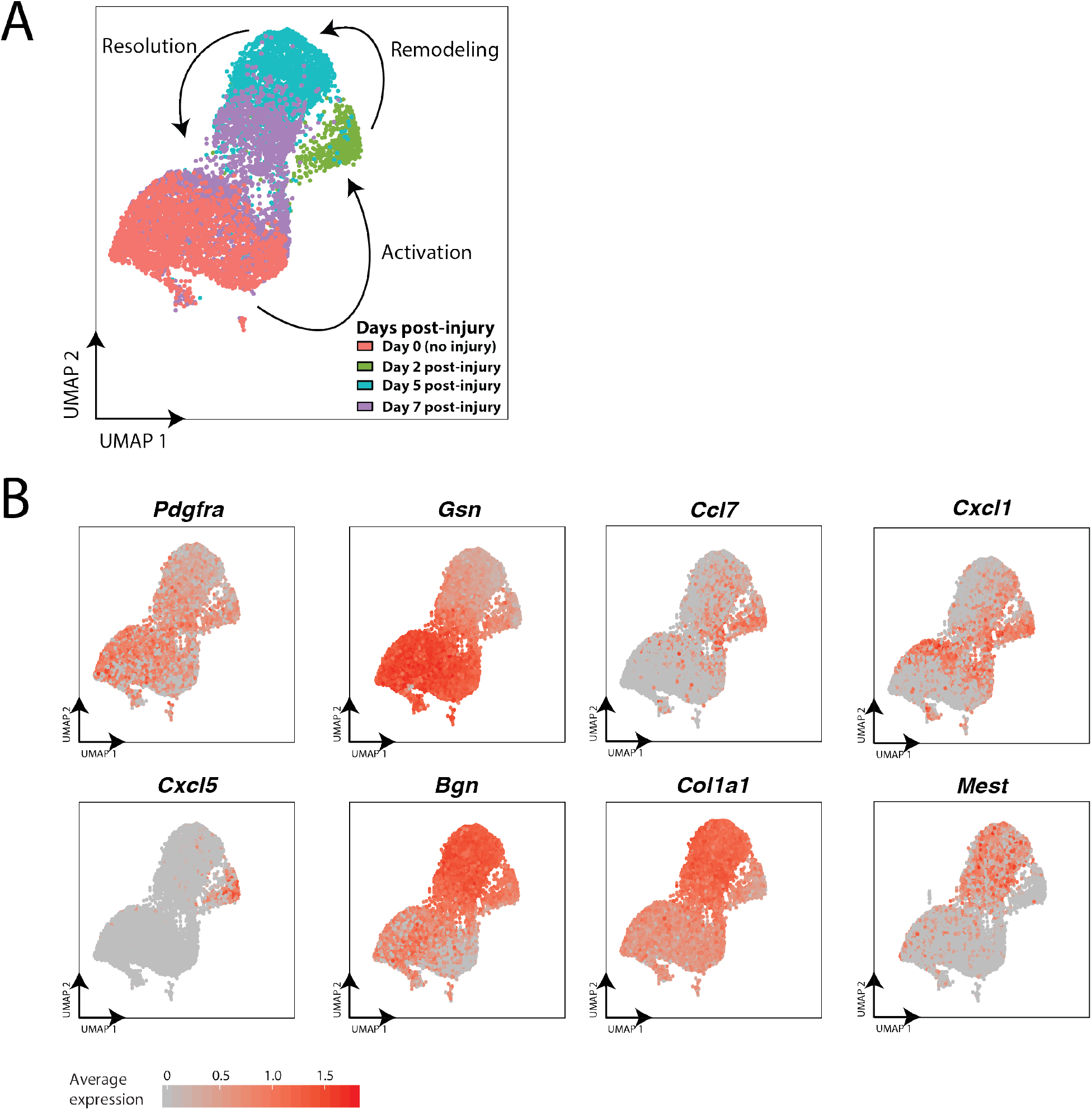
Gene expression dynamics of FAPs markers post-injury. (**A**) UMAP projection of single-FAP transcriptome colored by day post-injury. (**B**) Expression level of key FAP genes that describe the shift in gene expression profile post-injury.

**Figure S3.**
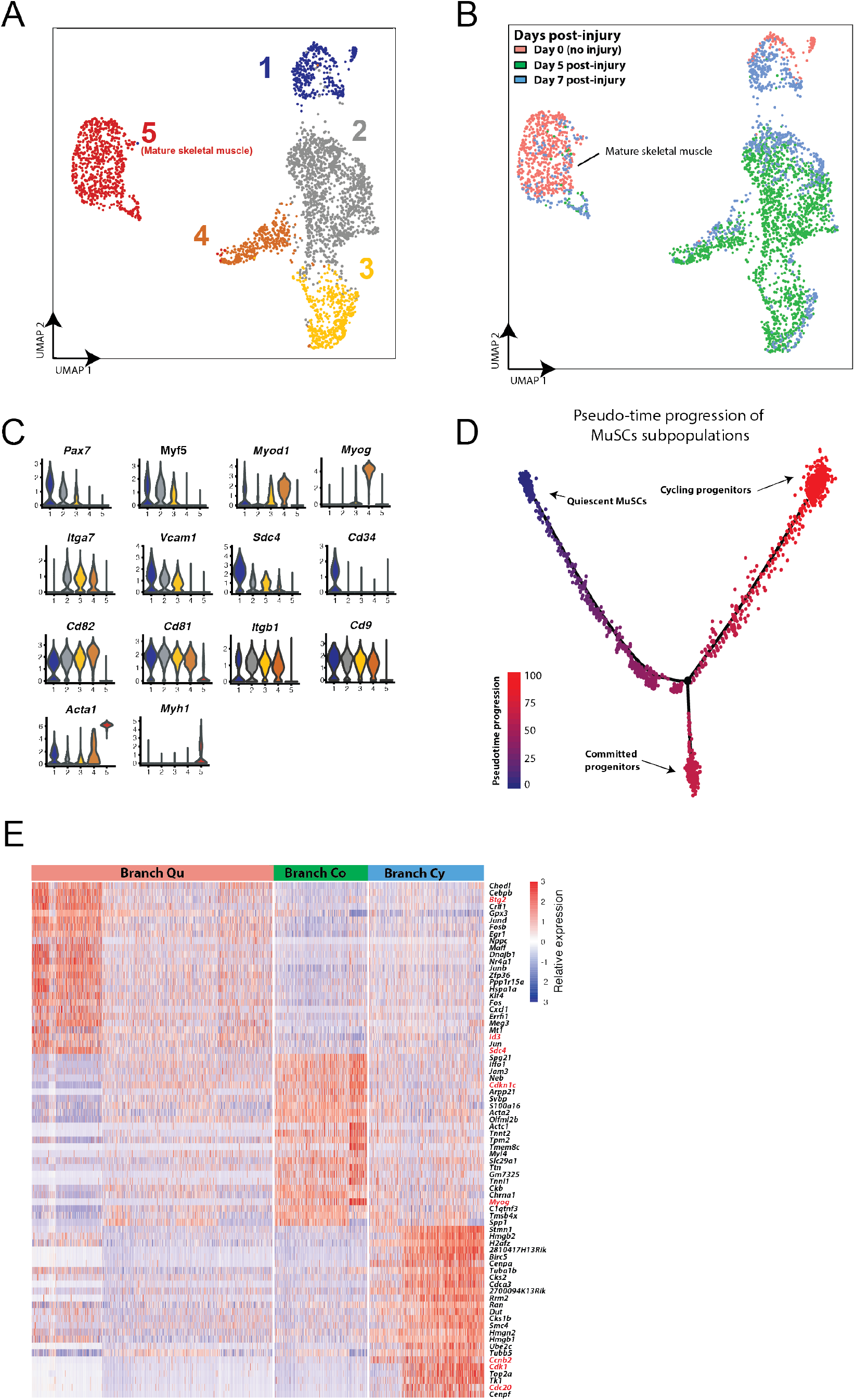
MuSCs heterogeneity in gene expression pre- and post-injury. (**A**) UMAP projection of MuSC and mature skeletal muscle single-cell transcriptomes (3,276 total cells from day 0, 5, 7) labeled by unsupervised SNN clustering. Clustering reveals 5 subpopulations of cells, including mature skeletal muscle cells (5) and MuSCs (1-4) that are composed of cells from different days post-injury. (**B**) UMAP projection of cells in (A) labeled by day post-injury. (**C**) Violin plots representing the expression level of key muscle genes in the subpopulations identified by SNN clustering (A). (**D**) Single-cell feature plot visualization of pseudo-time progression model of MuSC activation and commitment. (E) Top 75 genes differentially expressed in the three branches of the MuSC trajectory identified by Monocle (**Fig. 3B**) and ordered by branch and then pseudo-time progression.

**Figure S4.**
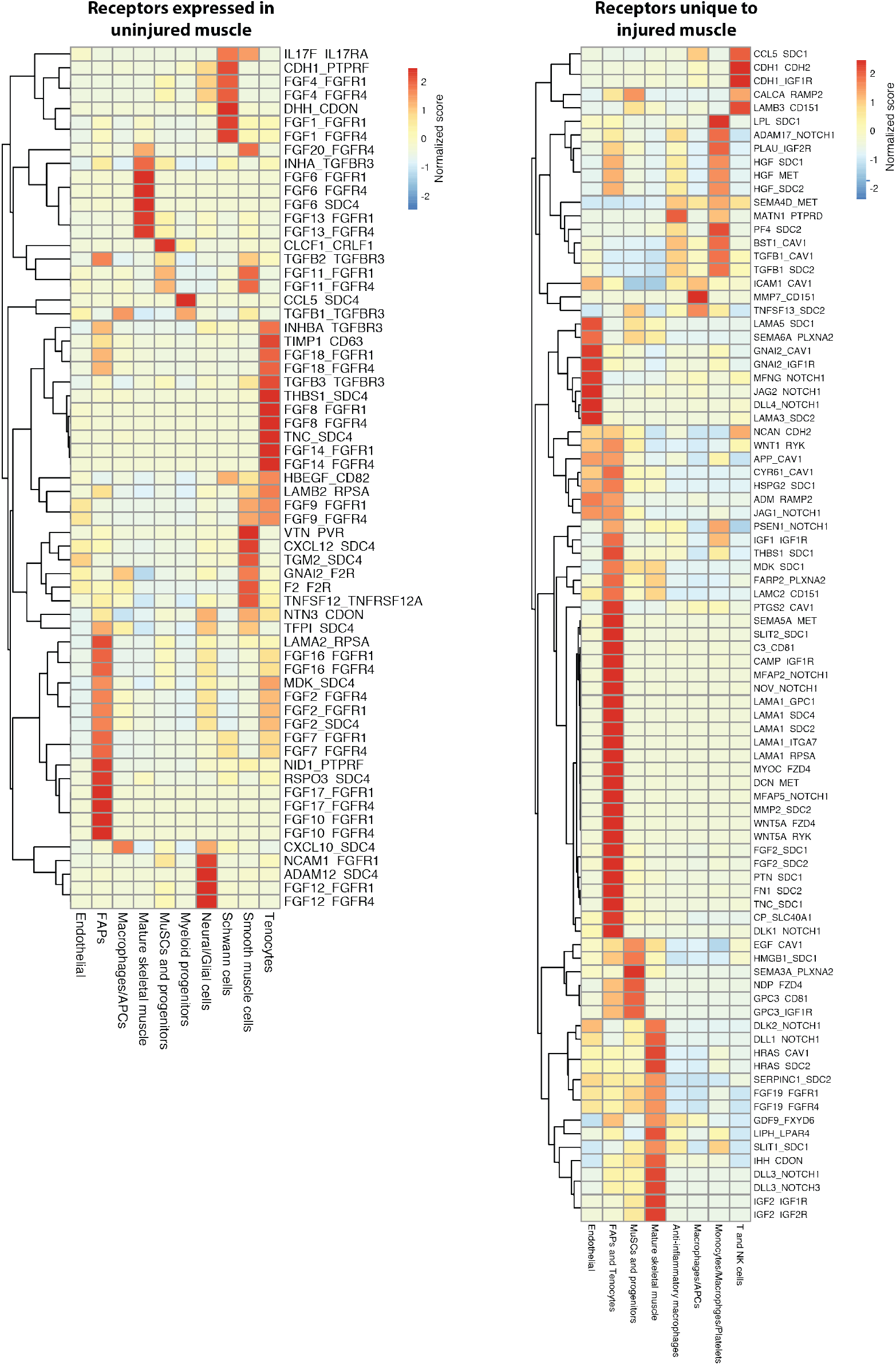
Ligand-receptor interaction score heatmap. The heatmap represents score between receptors differentially expressed in MuSCs and progenitor cells and ligands on other cell types. Columns represent the cell type expressing the ligand and includes MuSCs and progenitors for autocrine interactions. Rows represent the ligand-receptor pair in the format LIGAND_RECEPTOR. The score for each pair has been normalized across ligand cell types with a positive value indicating that the pair has a high score for a particular ligand and cell type compared to other cell types.

**Figure S5.**
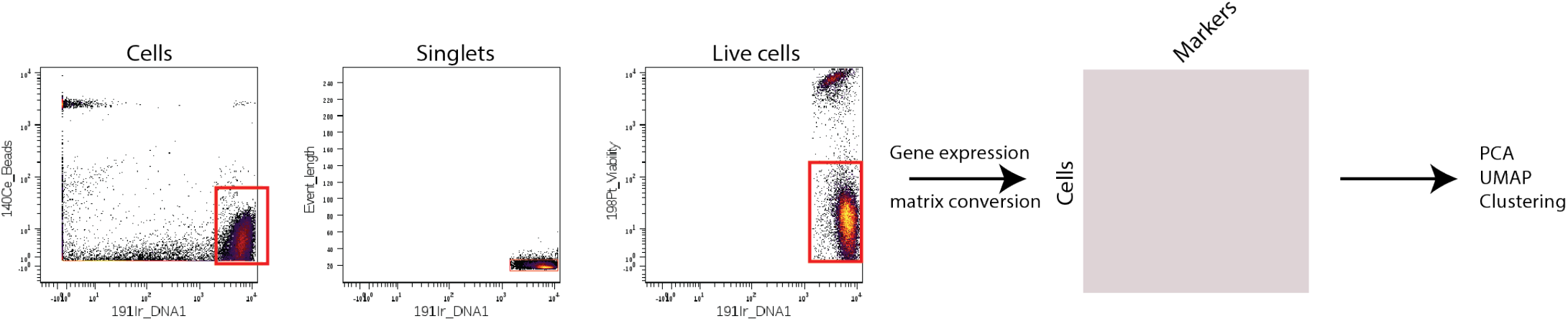
Mass cytometry (CyTOF) atlas of uninjured and injured muscle-tissue cells. Gating strategy before generation of a gene expression matrix. Cells were discriminated from debris using 140Ce beads, then from doublets by the event length, and finally from dead cells using a viability marker (198Pt-negative). The resulting gated FCS file was transformed into a gene expression matrix scaling by an inverse hyperbolic sine transformation.

**Table S1.**
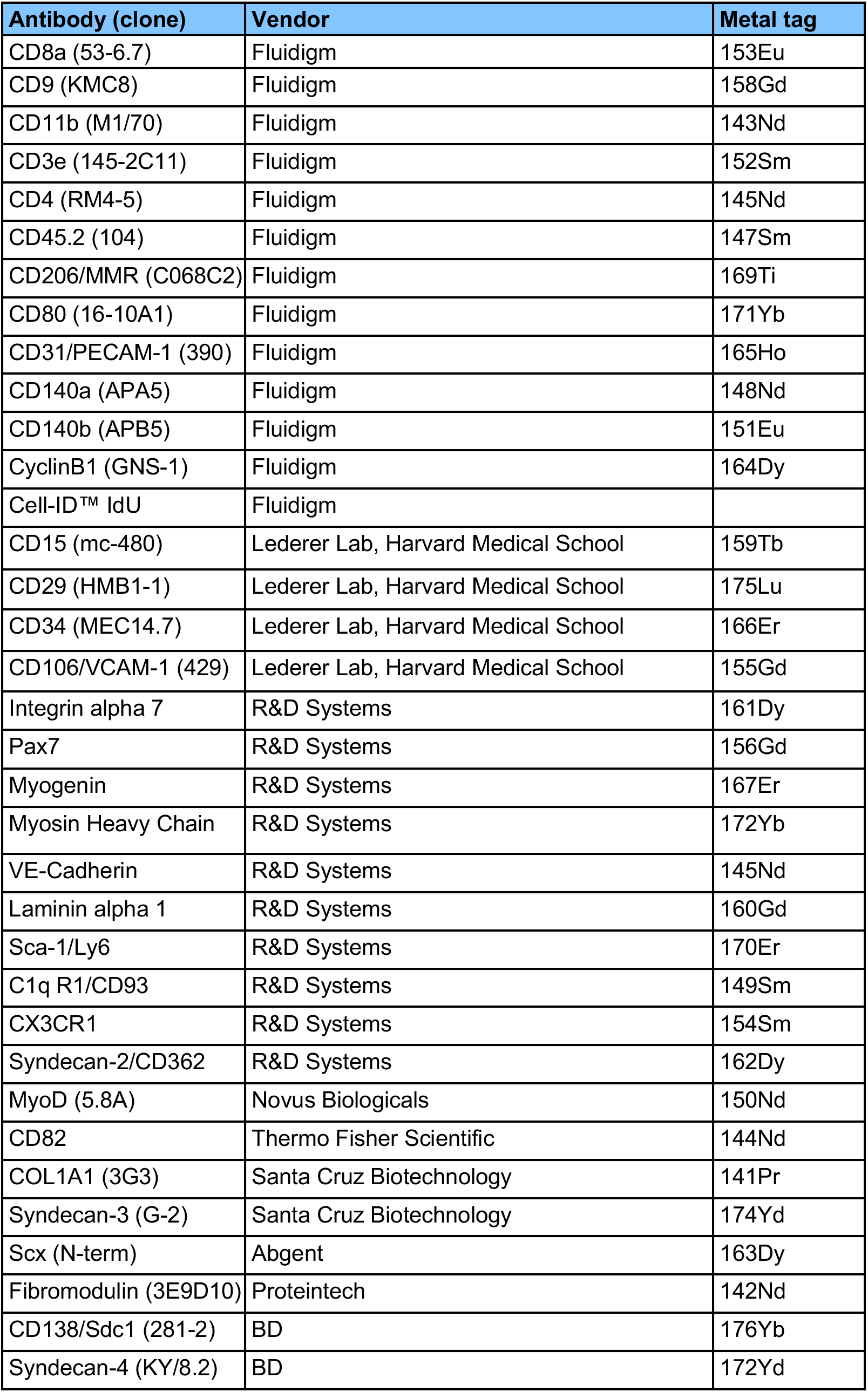
CyTOF reagent panel used for Figs 4 and S5.

## References

Baitsch, D., Bock, H.H., Engel, T., Telgmann, R., Muller-Tidow, C., Varga, G., Bot, M., Herz, J., Robenek, H., von Eckardstein, A., et al. (2011). Apolipoprotein E induces antiinflammatory phenotype in macrophages. Arterioscler Thromb Vasc Biol 31, 1160–1168.

Becht, E., McInnes, L., Healy, J., Dutertre, C.A., Kwok, I.W.H., Ng, L.G., Ginhoux, F., and Newell, E.W. (2018). Dimensionality reduction for visualizing single-cell data using UMAP. Nat Biotechnol.

Biressi, S., and Rando, T.A. (2010). Heterogeneity in the muscle satellite cell population. Semin Cell Dev Biol 21, 845–854.

Blau, H.M., Cosgrove, B.D., and Ho, A.T. (2015). The central role of muscle stem cells in regenerative failure with aging. Nat Med 21, 854–862.

Butler, A., Hoffman, P., Smibert, P., Papalexi, E., and Satija, R. (2018). Integrating single-cell transcriptomic data across different conditions, technologies, and species. Nat Biotechnol 36, 411–420.

Chakkalakal, J.V., Christensen, J., Xiang, W., Tierney, M.T., Boscolo, F.S., Sacco, A., and Brack, A.S. (2014). Early forming label-retaining muscle stem cells require p27kip1 for maintenance of the primitive state. Development 141, 1649–1659.

Chazaud, B. (2016). Inflammation during skeletal muscle regeneration and tissue remodeling: application to exercise-induced muscle damage management. Immunol Cell Biol 94, 140–145.

Chen, H., Lau, M.C., Wong, M.T., Newell, E.W., Poidinger, M., and Chen, J. (2016). Cytofkit: A Bioconductor Package for an Integrated Mass Cytometry Data Analysis Pipeline. PLoS Comput Biol 12, e1005112.

Cho, D.S., and Doles, J.D. (2017). Single cell transcriptome analysis of muscle satellite cells reveals widespread transcriptional heterogeneity. Gene 636, 54–63.

Christov, C., Chretien, F., Abou-Khalil, R., Bassez, G., Vallet, G., Authier, F.J., Bassaglia, Y., Shinin, V., Tajbakhsh, S., Chazaud, B., et al. (2007). Muscle satellite cells and endothelial cells: close neighbors and privileged partners. Mol Biol Cell 18, 1397–1409.

Cornelison, D.D., Wilcox-Adelman, S.A., Goetinck, P.F., Rauvala, H., Rapraeger, A.C., and Olwin, B.B. (2004). Essential and separable roles for Syndecan-3 and Syndecan-4 in skeletal muscle development and regeneration. Genes Dev 18, 2231–2236.

Cornelison, D.D., and Wold, B.J. (1997). Single-cell analysis of regulatory gene expression in quiescent and activated mouse skeletal muscle satellite cells. Dev Biol 191, 270–283.

Cosgrove, B.D., Gilbert, P.M., Porpiglia, E., Mourkioti, F., Lee, S.P., Corbel, S.Y., Llewellyn, M.E., Delp, S.L., and Blau, H.M. (2014). Rejuvenation of the muscle stem cell population restores strength to injured aged muscles. Nat Med 20, 255–264.

Dell’Orso, S., Juan, A.H., Ko, K.D., Naz, F., Perovanovic, J., Gutierrez-Cruz, G., Feng, X., and Sartorelli, V. (2019). Single cell analysis of adult mouse skeletal muscle stem cells in homeostatic and regenerative conditions. Development 146.

Docheva, D., Hunziker, E.B., Fassler, R., and Brandau, O. (2005). Tenomodulin is necessary for tenocyte proliferation and tendon maturation. Mol Cell Biol 25, 699–705.

Farioli-Vecchioli, S., Saraulli, D., Costanzi, M., Leonardi, L., Cina, I., Micheli, L., Nutini, M., Longone, P., Oh, S.P., Cestari, V., et al. (2009). Impaired terminal differentiation of hippocampal granule neurons and defective contextual memory in PC3/Tis21 knockout mice. PloS one 4, e8339.

Gaylinn, B.D., Eddinger, T.J., Martino, P.A., Monical, P.L., Hunt, D.F., and Murphy, R.A. (1989). Expression of nonmuscle myosin heavy and light chains in smooth muscle. Am J Physiol 257, C997–1004.

Giordani, L., He, G.J., Negroni, E., Sakai, H., Law, J.Y.C., Siu, M.M., Wan, R., Corneau, A., Tajbakhsh, S., Cheung, T.H., et al. (2019). High-Dimensional Single-Cell Cartography Reveals Novel Skeletal Muscle-Resident Cell Populations. Mol Cell 74, 609–621 e606.

Heredia, J.E., Mukundan, L., Chen, F.M., Mueller, A.A., Deo, R.C., Locksley, R.M., Rando, T.A., and Chawla, A. (2013). Type 2 innate signals stimulate fibro/adipogenic progenitors to facilitate muscle regeneration. Cell 153, 376–388.

Ho, M.M., Manughian-Peter, A., Spivia, W.R., Taylor, A., and Fraser, D.A. (2016). Macrophage molecular signaling and inflammatory responses during ingestion of atherogenic lipoproteins are modulated by complement protein C1q. Atherosclerosis 253, 38–46.

Hwang, B., Lee, J.H., and Bang, D. (2018). Single-cell RNA sequencing technologies and bioinformatics pipelines. Exp Mol Med 50, 96.

Janssen, E., Dzeja, P.P., Oerlemans, F., Simonetti, A.W., Heerschap, A., de Haan, A., Rush, P.S., Terjung, R.R., Wieringa, B., and Terzic, A. (2000). Adenylate kinase 1 gene deletion disrupts muscle energetic economy despite metabolic rearrangement. EMBO J 19, 6371–6381.

Joe, A.W., Yi, L., Natarajan, A., Le Grand, F., So, L., Wang, J., Rudnicki, M.A., and Rossi, F.M. (2010). Muscle injury activates resident fibro/adipogenic progenitors that facilitate myogenesis. Nat Cell Biol 12, 153–163.

Kotecha, N., Krutzik, P.O., and Irish, J.M. (2010). Web-based analysis and publication of flow cytometry experiments. Curr Protoc Cytom Chapter 10, Unit10 17.

Kuang, S., Kuroda, K., Le Grand, F., and Rudnicki, M.A. (2007). Asymmetric self-renewal and commitment of satellite stem cells in muscle. Cell 129, 999–1010.

Kumar, D., Shadrach, J.L., Wagers, A.J., and Lassar, A.B. (2009). Id3 is a direct transcriptional target of Pax7 in quiescent satellite cells. Mol Biol Cell 20, 3170–3177.

Kuschel, R., Deininger, M.H., Meyermann, R., Bornemann, A., Yablonka-Reuveni, Z., and Schluesener, H.J. (2000). Allograft inflammatory factor-1 is expressed by macrophages in injured skeletal muscle and abrogates proliferation and differentiation of satellite cells. J Neuropathol Exp Neurol 59, 323–332.

Liu, Z., Jin, Y.Q., Chen, L., Wang, Y., Yang, X., Cheng, J., Wu, W., Qi, Z., and Shen, Z. (2015). Specific marker expression and cell state of Schwann cells during culture in vitro. PloS one 10, e0123278.

Lukjanenko, L., Jung, M.J., Hegde, N., Perruisseau-Carrier, C., Migliavacca, E., Rozo, M., Karaz, S., Jacot, G., Schmidt, M., Li, L., et al. (2016). Loss of fibronectin from the aged stem cell niche affects the regenerative capacity of skeletal muscle in mice. Nat Med 22, 897–905.

Lyons, G.E., Ontell, M., Cox, R., Sassoon, D., and Buckingham, M. (1990). The expression of myosin genes in developing skeletal muscle in the mouse embryo. J Cell Biol 111, 1465–1476.

Machado, L., Esteves de Lima, J., Fabre, O., Proux, C., Legendre, R., Szegedi, A., Varet, H., Ingerslev, L.R., Barres, R., Relaix, F., et al. (2017). In Situ Fixation Redefines Quiescence and Early Activation of Skeletal Muscle Stem Cells. Cell reports 21, 1982–1993.

Motohashi, N., and Asakura, A. (2014). Muscle satellite cell heterogeneity and self-renewal. Frontiers in cell and developmental biology 2.

Novak, M.L., and Koh, T.J. (2013). Macrophage phenotypes during tissue repair. J Leukoc Biol 93, 875–881.

Pawlikowski, B., Orion Vogler, T., Gadek, K., and Olwin, B. (2017). Regulation of Skeletal Muscle Stem Cells by Fibroblast Growth Factors. Dev Dyn.

Pisconti, A., Banks, G.B., Babaeijandaghi, F., Betta, N.D., Rossi, F.M., Chamberlain, J.S., and Olwin, B. B. (2016). Loss of niche-satellite cell interactions in syndecan-3 null mice alters muscle progenitor cell homeostasis improving muscle regeneration. Skelet Muscle 6, 34.

Pisconti, A., Bernet, J.D., and Olwin, B.B. (2012). Syndecans in skeletal muscle development, regeneration and homeostasis. Muscles Ligaments Tendons J 2, 1–9.

Pisconti, A., Cornelison, D.D., Olguin, H.C., Antwine, T.L., and Olwin, B.B. (2010). Syndecan-3 and Notch cooperate in regulating adult myogenesis. J Cell Biol 190, 427–441.

Porpiglia, E., Samusik, N., Ho, A.T., Cosgrove, B.D., Mai, T., Davis, K.L., Jager, A., Nolan, G.P., Bendall, S.C., Fantl, W.J., et al. (2017). High-resolution myogenic lineage mapping by single-cell mass cytometry. Nat Cell Biol 19, 558–567.

Qiu, X., Mao, Q., Tang, Y., Wang, L., Chawla, R., Pliner, H.A., and Trapnell, C. (2017). Reversed graph embedding resolves complex single-cell trajectories. Nature methods 14, 979–982.

Ren, H., Yin, P., and Duan, C. (2008). IGFBP-5 regulates muscle cell differentiation by binding to IGF-II and switching on the IGF-II auto-regulation loop. J Cell Biol 182, 979–991.

Rocheteau, P., Gayraud-Morel, B., Siegl-Cachedenier, I., Blasco, M.A., and Tajbakhsh, S. (2012). A subpopulation of adult skeletal muscle stem cells retains all template DNA strands after cell division. Cell 148, 112–125.

Sacco, A., Doyonnas, R., Kraft, P., Vitorovic, S., and Blau, H.M. (2008). Self-renewal and expansion of single transplanted muscle stem cells. Nature 456, 502–506.

Schweitzer, R., Chyung, J.H., Murtaugh, L.C., Brent, A.E., Rosen, V., Olson, E.N., Lassar, A., and Tabin, C.J. (2001). Analysis of the tendon cell fate using Scleraxis, a specific marker for tendons and ligaments. Development 128, 3855–3866.

Skelly, D.A., Squiers, G.T., McLellan, M.A., Bolisetty, M.T., Robson, P., Rosenthal, N.A., and Pinto, A.R. (2018). Single-Cell Transcriptional Profiling Reveals Cellular Diversity and Intercommunication in the Mouse Heart. Cell reports 22, 600–610.

Sousa-Victor, P., Gutarra, S., Garcia-Prat, L., Rodriguez-Ubreva, J., Ortet, L., Ruiz-Bonilla, V., Jardi, M., Ballestar, E., Gonzalez, S., Serrano, A.L., et al. (2014). Geriatric muscle stem cells switch reversible quiescence into senescence. Nature 506, 316–321.

Stuart, T., and Satija, R. (2019). Integrative single-cell analysis. Nat Rev Genet 20, 257–272.

Tidball, J.G. (2017). Regulation of muscle growth and regeneration by the immune system. Nat Rev Immunol 17, 165–178.

Tierney, M.T., and Sacco, A. (2016). Satellite Cell Heterogeneity in Skeletal Muscle Homeostasis. Trends Cell Biol 26, 434–444.

Tierney, M.T., Stec, M.J., Rulands, S., Simons, B.D., and Sacco, A. (2018). Muscle Stem Cells Exhibit Distinct Clonal Dynamics in Response to Tissue Repair and Homeostatic Aging. Cell Stem Cell 22, 119–127 e113.

Uezumi, A., Ito, T., Morikawa, D., Shimizu, N., Yoneda, T., Segawa, M., Yamaguchi, M., Ogawa, R., Matev, M.M., Miyagoe-Suzuki, Y., et al. (2011). Fibrosis and adipogenesis originate from a common mesenchymal progenitor in skeletal muscle. J Cell Sci 124, 3654–3664.

van Velthoven, C.T.J., de Morree, A., Egner, I.M., Brett, J.O., and Rando, T.A. (2017). Transcriptional Profiling of Quiescent Muscle Stem Cells In Vivo. Cell reports 21, 1994–2004.

Wagner, A., Regev, A., and Yosef, N. (2016). Revealing the vectors of cellular identity with single-cell genomics. Nat Biotechnol 34, 1145–1160.

Wang, Y.X., and Rudnicki, M.A. (2011). Satellite cells, the engines of muscle repair. Nat Rev Mol Cell Biol 13, 127–133.

Wosczyna, M.N., and Rando, T.A. (2018). A Muscle Stem Cell Support Group: Coordinated Cellular Responses in Muscle Regeneration. Dev Cell 46, 135–143.

Yin, H., Price, F., and Rudnicki, M.A. (2013). Satellite cells and the muscle stem cell niche. Physiol Rev 93, 23–67.

Yuniati, L., Scheijen, B., van der Meer, L.T., and van Leeuwen, F.N. (2019). Tumor suppressors BTG1 and BTG2: Beyond growth control. J Cell Physiol 234, 5379–5389.

Zordan, P., Rigamonti, E., Freudenberg, K., Conti, V., Azzoni, E., Rovere-Querini, P., and Brunelli, S. (2014). Macrophages commit postnatal endothelium-derived progenitors to angiogenesis and restrict endothelial to mesenchymal transition during muscle regeneration. Cell Death Dis 5, e1031.

